# L-Phenylalanine is a metabolic checkpoint of human Th2 cells

**DOI:** 10.1101/2024.07.29.605231

**Authors:** Abhijeet J Kulkarni, Juan Rodriguez-Coira, Nino Stocker, Urszula Radzikowska, Antonio J García-Cívico, María Isabel Delgado Dolset, Nuria Contreras, Inés Jardón Parages, Vanessa Saiz Sanchez, Pilar Serrano, Elena Izquierdo, Cristina Gomez-Casado, Javier Sanchez-Solares, Carmela Pablo-Torres, David Obeso, Carmen Moreno-Aguilar, Maria Luisa Espinazo, Andrzej Eljaszewicz, Jana Koch, Katja Baerenfaller, Anja Heider, Ge Tan, Maria M Escribese, Berta Ruiz-Leon, Cezmi A Akdis, Rafael J. Argüello, Domingo Barber, Alma Villaseñor, Milena Sokolowska

**Author notes:** **Corresponding authors:** Milena Sokolowska (Lead corresponding author) –; 0Alma Villaseñor –. Equal Contribution (Abhijeet J Kulkarni, Juan Rodriguez-Coira). Equal Senior Contribution and Corresponding Authors (Alma Villaseñor, Milena Sokolowska).

## Abstract

After the primary response, circulating memory CD4^+^T effector and T regulatory cells (Treg) regulate recall responses, which are impaired in allergy. Using mass spectrometry, we discovered distinct metabolomes of these cells in humans and their unique enrichment in amino acids. By assessing energy metabolism in *in vitro* and *ex vivo* single-cell analyses, we determined that increased intracellular L-phenylalanine boosts glycolysis while limiting OXPHOS in CD4^+^T, memory CD4^+^T and Th2, but not in Treg cells. L-phenylalanine also restrains memory CD4^+^T proliferation in an IL4I1-dependent manner and inhibits Th2 cell proliferation and differentiation. RNA-sequencing, metabolomics, flow cytometry and proteomics, validated *in vitro* and across patients’ cohorts, revealed an impairment in LAT1-dependent transport of L-phenylalanine into Th2 cells in allergy with an increase in its intracellular processing, accompanied by an expansion of pathogenic Th2 cells. Thus, our study identifies L-phenylalanine as a checkpoint in the development, energy metabolism and function of Th2 cells.

## Introduction

Upon activation, naïve CD4^+^T cells clonally expand and differentiate into several specialized effector (Teff) subsets, including T helper (Th) 1, Th2 and Th17, depending on the nature of antigen challenge and cues from the microenvironment.^1^ Regulatory CD4^+^CD25^+^T cells (Treg) arise either from the thymus or from peripheral conversion of conventional T cells.^2,3^ After initial clonal expansion, majority of CD4^+^Teff cells are quickly removed in the absence of antigen, but a fraction of them form long-lived memory T cells, persisting in tissues, lymphatic organs, and, importantly, in circulation.^1,4,5^ Memory CD4^+^T cell subsets are able to mount quicker and greater recall Teff and Treg response after subsequent encounter with the antigen.^1^ Quiescent naïve and memory CD4^+^T cells, and Treg cells tend to primarily use fatty acid oxidation (FAO) and oxidative phosphorylation (OXPHOS)^6–9^, although, in human Treg cells, glycolysis and fatty acid metabolism are also very active and needed for their suppressive functions.^10,11^ Post antigen encounter, co-stimulation, and signals from cytokines, CD4^+^Teff and Treg subsets undergo further changes in their glucose, amino acid and fatty acid metabolism fueling subset-specific proliferation and function.^7,12–16^ Maintaining the equilibrium among memory CD4^+^Teff and Treg subsets is essential for homeostasis and prevention of inflammatory or autoimmune diseases. However, while the metabolic requirements of naïve CD4^+^T cells upon initial activation and differentiation into the effector and regulatory subsets is well studied^17,18^, much less is known about the metabolic programs of human circulating memory CD4^+^Teff and Treg cells.

T cell function and metabolism are controlled by amino acids via acquisition from the extracellular milieu through cellular transporters, intracellular storage, and recycling within different metabolic pathways, especially glycolysis, citric acid (TCA) cycle and OXPHOS.^13,19–21^ Expression of L-Leucine (Leu), L-Isoleucine (Ile), L-Tryptophan and other large neutral amino acids transporters such as LAT1 (*SLC7A5*), LAT2 (*SLC7A8*), CD98 (*SLC3A2*), as well as *SLC1A5, SLC38A2, SLC7A1* and others transporting L-Glutamine, L-Serine or L-Arginine (Arg), increases during activation of CD4^+^ and CD8^+^T cells.^13,22^ Expression of these transporters, sensing extracellular amino acids in a mechanistic target of rapamycin (mTOR)-dependent way, as well as their intracellular utilization, regulate CD4^+^Teff and Treg cells metabolic reprogramming and subsequently their proliferation, differentiation and function.^13,20–24^ Amino acid metabolizing enzymes, such as Interleukin 4 Induced Gene 1 (IL4I1)^25–28^ or glutamate oxaloacetate transaminase 1 (GOT1)^29^, also affect Teff and Treg cells. L-Phenylalanine (Phe), also transported by the same set of transporters might be partly metabolised by the same enzymes, but its role in metabolism, proliferation, and differentiation of memory CD4^+^T cells has not been explored.

Allergic diseases such as allergic asthma, food allergy or rhinitis are common type 2 inflammatory diseases, characterized by skewed Teff-Treg responses, where pathogenic Th2 cells cannot be efficiently suppressed by impaired Treg cells. In humans, pathogenic CRTH2^+^ or CD161^+^Th2 cells, overexpressing GATA3 and PPARγ, producing IL-4, IL-5, IL-9, and IL-13, are predominantly involved in the onset and maintenance phase of the disease.^30–34^ Th2 cells require mTORC1 for engagement of glycolysis and cell cycle entry^35^, mTORC2 for cell survival and migratory functions^23,36^, and fatty acid synthesis (FAS) for lineage development.^37^ We and others demonstrated that in severe forms of allergic diseases there are systemic and cellular alterations in amino acids, fatty acids, and glycolysis.^38–41^ In patients with severe allergic asthma many amino acids including arginine, phenylalanine, taurine, and xanthine are increased in plasma^42^, but how it affects metabolism and functions of memory Th2 and Treg cells remain elusive.

Here, we report that Phe constitutes a previously unknown checkpoint in the development, energy metabolism and function of Th2 cells. Using high-resolution mass spectrometry, we discovered that metabolome profiles of *ex vivo* sorted circulating human memory CD4^+^Teff and Treg cells are enriched in amino acid metabolites and clearly differentiate these cell types. Combining *in vitro* functional experiments and knockdown approaches in primary human CD4^+^T cells, memory CD4^+^T cells and *in vitro* differentiated Th2 cells with *ex vivo* single cell energy metabolism profiling, we discovered that high level of intracellular Phe increases glycolysis, but restricts OXPHOS, as well as affects memory CD4^+^T cell proliferation and Th2 differentiation program in IL4I1-dependent manner. Finally, analyzing five different cohorts of allergic patients by combining RNA-seq analysis, metabolomics, proteomics, and flow cytometry, we noted that in allergy there is an impairment of SLC7A5-dependent transport of Phe into Th2 cells and an increase in its intracellular processing, leading to an excessive expansion of pathogenic Th2 cells.

## Results

### Circulating human memory CD4^+^T effector and regulatory T cells have different metabolic profiles and are enriched in amino acids

We first aimed to assess the metabolome of circulating human memory T cells in an unbiased manner after *ex vivo* cell sorting without any culturing steps in healthy subjects (n=6). We focused on analyzing the memory T effector (Teff) cells (CD3^+^CD4^+^CD8^-^CD45RA^-^ CD127^+^CD25^-^) and memory T regulatory (Treg) cells (CD3^+^CD4^+^CD8^-^CD45RA^-^CD127^-^ CD25^+^) by untargeted metabolomics and lipidomics **(Figure 1A, 1B; Figure S1, Table S1)**. Following data processing and analysis, we obtained a total of 195 and 233 metabolites for Teff and Treg cells, respectively, of which 133 (45%) were shared by both cell types **(Figure 1C; Table S2)**. Principal Component Analysis (PCA; **Figure 1D**) and orthogonal partial-least discriminant analysis (OPLS-DA; **Figure 1E**) models revealed a complete separation between these populations even though their metabolites belonged to similar major families (**Figure 1F**). Carboxylic acids and their derivatives, which include major amino acids such as Phe, Arg, and histidine, represented the largest group in both cell types (>20%, **Figure 1F**). Fatty acyls, glycerophospholipids and organooxygen compounds (including glucose) were the other major biochemical classes in both cell types **(Table S3)**. The metabolomes of both cell subsets were enriched in metabolites related to pathways such as amino acids biosynthesis and metabolism, vitamin synthesis as well as in energy metabolism (glycolysis/gluconeogenesis, mTOR signalling). Moreover, Teff cells were uniquely enriched in glycerophospholipid metabolism pathway, while Treg metabolome was solely over-represented in fatty acid biosynthesis **(Figure 1G, Tables S4, S5)**.

**Figure 1.**
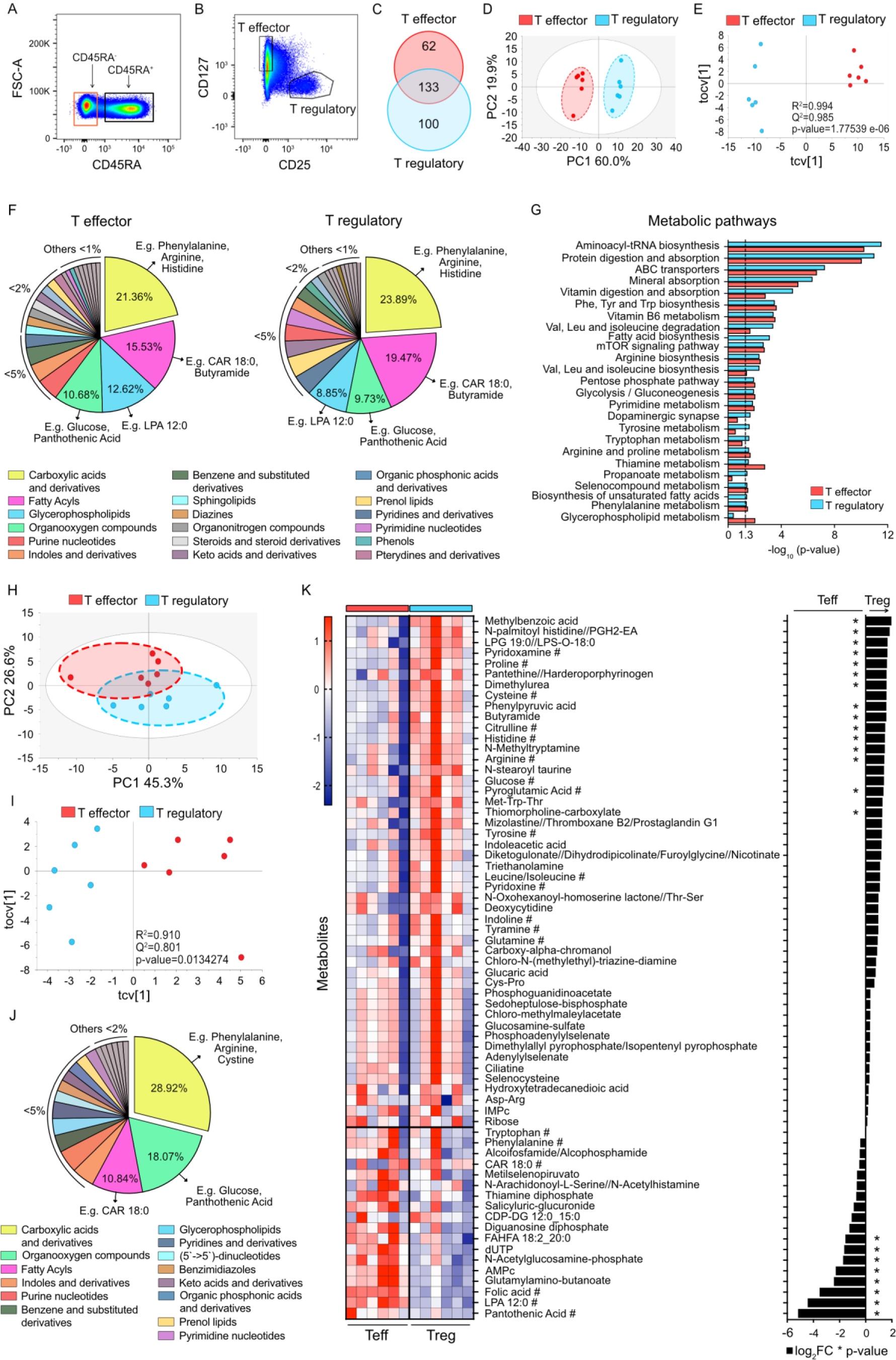
Metabolomics of circulating human memory CD4^+^T effector and T regulatory cells reveals distinct metabolic profiles, enriched in amino acid metabolic pathways. (**A-B**) Intermediate gate depicting memory (CD45RA^-^) and naive (CD45RA^+^) CD3^+^CD4^+^T cells (**A**) and final sorting gate (**B**) of circulating memory CD4^+^T effector (CD3^+^CD4^+^CD8^-^CD45RA^-^ CD127^+^CD25^-^) and T regulatory (CD3^+^CD4^+^CD8^-^CD45RA^-^CD127^-^CD25^+^) cells. Figure S1 depicts the complete gating strategy. (**C**) Venn diagram representing intracellular metabolites (n=295) detected by untargeted mass-spectrometry metabolomics and lipidomics in memory CD4^+^Teff (red) and Treg (blue) cells from healthy individuals. (**D-E**) Principal Component Analysis (PCA) (**D**) and Orthogonal Partial-Least Discriminant Analysis (OPLS-DA) (**E**) models revealing separation of memory CD4^+^Teff and Treg cells, based on all (shared and unique) metabolites. Data was logarithmic-transformed and pareto-scaled (log x Par). (**F**) Pie chart representing biochemical composition of all (shared and unique) metabolites detected in Teff and Treg cells, ordered by abundance. (**G**) Metabolic pathways analysis in memory CD4^+^Teff and Treg cells. Significant (-log_10_(p-value)>1.3) and corresponding pathways (n=25) are shown for either Teff, Treg, or both. Over-representation analysis was performed by Integrated Molecular Pathway Level Analysis (IMPaLA)^99^ including shared and unique metabolites. (**H-I**) PCA (**H**) and OPLS-DA (**I**) models of memory CD4^+^Teff and Treg cells revealing separation based on shared metabolites only (n=133). Data was log x Par. (**J**) Pie chart representing biochemical composition of shared metabolites in memory CD4^+^Teff and Treg cells, ordered by abundance. (**K**) Heatmap (left) and bar graph of fold changes (Log_2_FC) (right) of identified shared metabolites in memory CD4^+^Teff vs Treg cells. Data was logarithmic-transformed and unit variance-scaled. *p<0.05. Multiplicated metabolites result from detection in different analyses (from different extraction methods), measurement in positive and negative ionization modes, and/or found clustered with different adducts. # indicates metabolites with better score for their identity. (A-K) Analysis done in n=6 different healthy donors. (F and J) Metabolites are organised in main biochemical classes according to Human Metabolome Database (HMDB v. 2022)^100^. Examples of metabolites are shown in major classes (> 5%).

Since the strongest enrichment in both cell types was present due to the highly abundant shared metabolites, we focused primarily on these compounds. Interestingly, we found that shared metabolites can also clearly separate Teff cells from Treg cells in PCA and OPLS-DA models **(Figure 1H-1I)**, suggesting different abundance of these metabolites. Regarding the biochemical classes of shared metabolites, likewise, carboxylic acids and their derivatives class, including many amino acids, organooxygen compounds and fatty acyls represented the biggest fractions **(Figure 1J, Table S6)**. Among these, we found some organooxygen compounds, such as pantothenic acid, and glycerophospholipids, such as lysophosphatidic acid (LPA), to be significantly elevated in Teff cells, while fatty acyls like N-palmitoyl histidine//PGH2-EA and carboxylic acids and derivatives, such as Arg, histidine and citrulline, were significantly higher in Treg cells **(Figure 1K, Table S7)**.

In summary, we found here that the major similarities in metabolomes of *ex vivo* circulating human memory Teff and Treg cells include various amino acids and their derivatives. Interestingly, however, the abundances of several of these compounds significantly differ between these two cell types, suggesting differential usage or activation of different metabolic pathways.

### L-Phenylalanine increases glycolysis and inhibits oxidative phosphorylation (OXPHOS) in CD4^+^T cells

Since we found that amino acids constituted the largest group of total and shared metabolites, and we noticed enrichment in energy metabolism pathways shared between memory CD4^+^Teff and Treg cells, we analysed the effects of some of these amino acids on glycolysis and OXPHOS in human primary CD4^+^T cells. There are reports on the role of Arg in human T cell metabolism^19,20^, whereas very little is known in terms of Phe; thus, we first aimed to assess their effects on energy metabolism of human primary CD4^+^T cells at baseline and after rapid polyclonal activation to emulate TCR recognition. As expected, activated total CD4^+^T cells displayed higher levels of inducible and compensatory glycolysis, and these parameters were further elevated after 72h incubation with increasing concentrations of Arg **(Figure 2A)**. We also observed increased levels of maximum respiratory capacity with higher concentrations of Arg **(Figure 2B)**. Similarly, activated total CD4^+^T cells incubated in additional 0.1mM and 1mM Phe for 72h had higher inducible and compensatory glycolysis **(Figure 2C).** Interestingly, the maximum respiratory capacity of cells incubated in additional 1mM of Phe was significantly lower compared to that of cells incubated in 0.1mM Phe supplemented media at this time point **(Figure 2D)** suggesting that higher levels of Phe negatively influence OXPHOS. When the same experimental setup was repeated in memory CD4^+^T cells treated with additional 0.1mM and 1mM of Phe, we observed again increased inducible and compensatory glycolysis with increasing doses of Phe. Likewise, the maximum respiratory capacity was again reduced in memory CD4^+^T cells incubated in 1mM Phe supplemented media as observed for total CD4^+^T cells **(Figure 2E)**.

**Figure 2.**
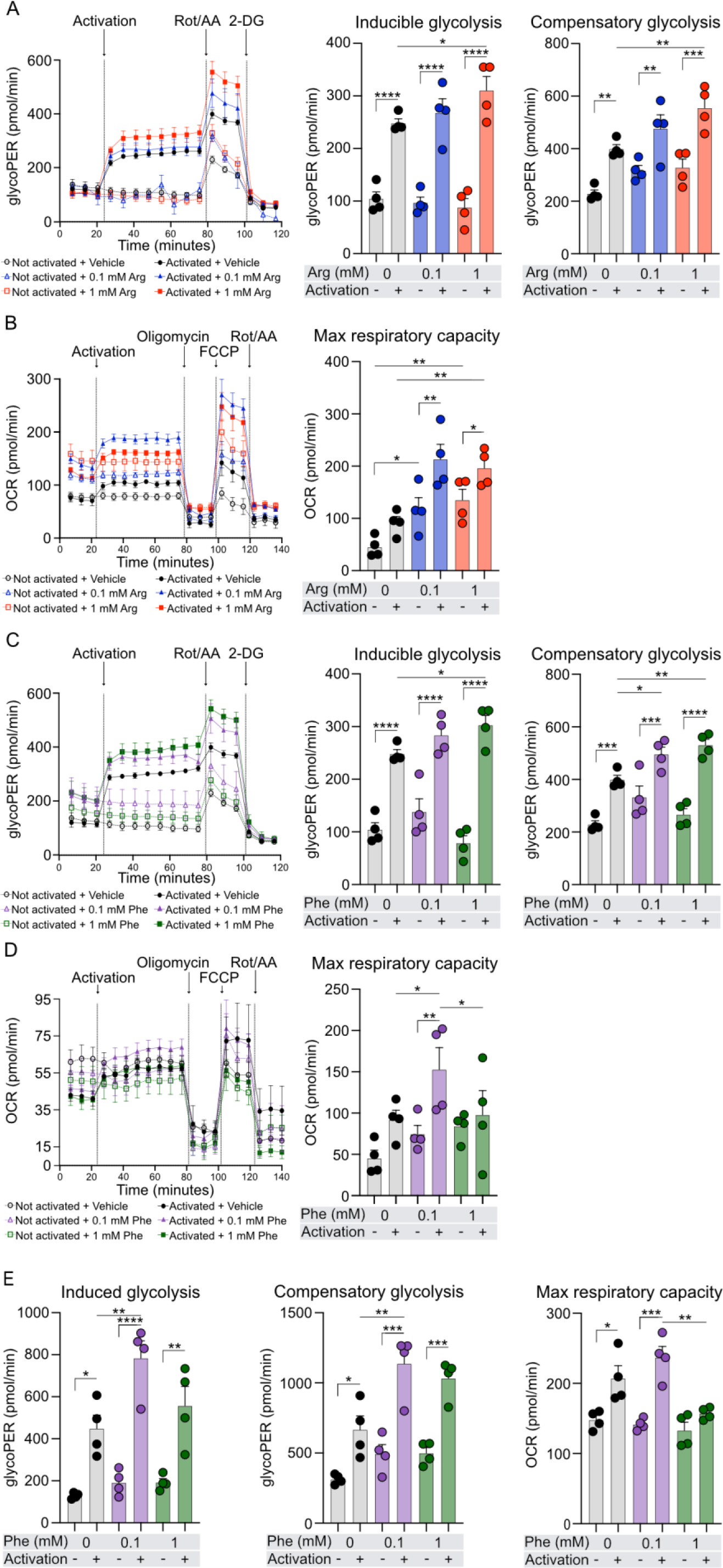
High level of L-phenylalanine enhances activation-induced glycolysis but inhibits OXPHOS, while arginine enhances activation-induced glycolysis and OXPHOS in human CD4^+^T and memory CD4^+^T cells. (**A**) Representative glycolytic proton efflux rate (glycoPER) graph of Seahorse Glycolytic Rate assay (left); quantification of inducible and compensatory glycolysis (right), of CD4^+^T cells treated in full medium with 0.1mM (blue), 1mM (red) Arg supplementation or Vehicle (grey) for 72h with/without acute CD2, CD3 and CD28 activation. (**B**) Representative oxygen consumption rate (OCR) graph of Seahorse Mito Stress test from one donor (left); quantification of maximum respiratory capacity (right) of CD4^+^T cells treated with Arg as in (A). (**C**) Representative glycoPER graph of Seahorse Glycolytic Rate assay (left); quantification of inducible and compensatory glycolysis (right) of CD4^+^T cells treated in full medium with Phe supplemented at concentrations of 0.1mM (violet), 1mM (green), or Vehicle (grey) for 72h, with/without acute CD2, CD3 and CD28 activation. (**D**) Representative OCR graph of Seahorse Mito Stress test (left); quantification of maximum respiratory capacity (right) of CD4^+^T cells treated with Phe as in (C) (**E**) Quantification of induced glycolysis, compensatory glycolysis, and maximum respiratory capacity of memory CD4^+^T cells treated with Phe as in (C and D). (A-D) Data are representative of three independent experiments in three different donors or (E) in one donor. Data were analyzed by One-way ANOVA with Fisher LSD test. Bar graphs represent mean±SEM. *p<0.05, **p<0.01, ***p<0.001, ****p<0.0001. Legend: Arg: L-arginine; Phe: L-phenylalanine; Rot/AA: rotenone/antimycin A; 2DG: 2-deoxyglucose; FCCP: carbonyl cyanide-p-trifluoromethoxyphenylhydrazone. All Seahorse measurements were normalized to total protein concentration in each well.

To sum up, here, we observed that while Arg induces both glycolysis and OXPHOS in total human CD4^+^T cells, Phe enhances glycolysis but impairs OXPHOS at higher doses in total and memory human CD4^+^T cells.

### L-Phenylalanine controls CD4^+^T cell proliferation via induction of Interleukin-4 induced 1 (IL4I1) enzyme

Considering the ability of Phe to differentially influence glycolysis and OXPHOS in total and memory CD4^+^T cells upon activation, we sought to determine if Phe also affects their ability to proliferate. Memory CD4^+^T cells incubated in full media supplemented additionally with 1mM Phe showed reduced proliferation after 72h and 120h of incubation **(Figure 3A)** with no significant difference in the number of dead cells **(Figure 3B)**. To gain insight into the mechanism behind this phenomenon, we assessed the mRNA expression of the Interleukin 4 induced gene 1 enzyme, *IL4I1*, that has been reported to limit effector T cell proliferation.^28,43,44^ We found that supplementation of CD4^+^T cells with high levels of Phe significantly upregulated mRNA expression of *IL4I1* **(Figure 3C).** Importantly, lower levels of Phe, which did not affect OXPHOS or proliferation of CD4^+^T cells, also did not influence *IL4I1* expression. To confirm the role of IL4I1 in blocking memory CD4^+^T cell proliferation, we carried out siRNA-based IL4I1 knockdown in memory CD4^+^T cells and assessed their proliferation. We observed significant knockdown of IL4I1 in these donors based on *IL4I1* qRT-PCR and Western blot analysis **(Figure 3D)**. We found a significant increase in memory CD4^+^T cells proliferation when IL4I1 was knocked down and cells were either activated with beads **(Figure 3E)** or non-activated **(Figure S2A)**. Importantly, IL4I1 knockdown did not affect cell viability as compared to control siRNA **(Figure 3F, Figure S2B)**.

**Figure 3.**
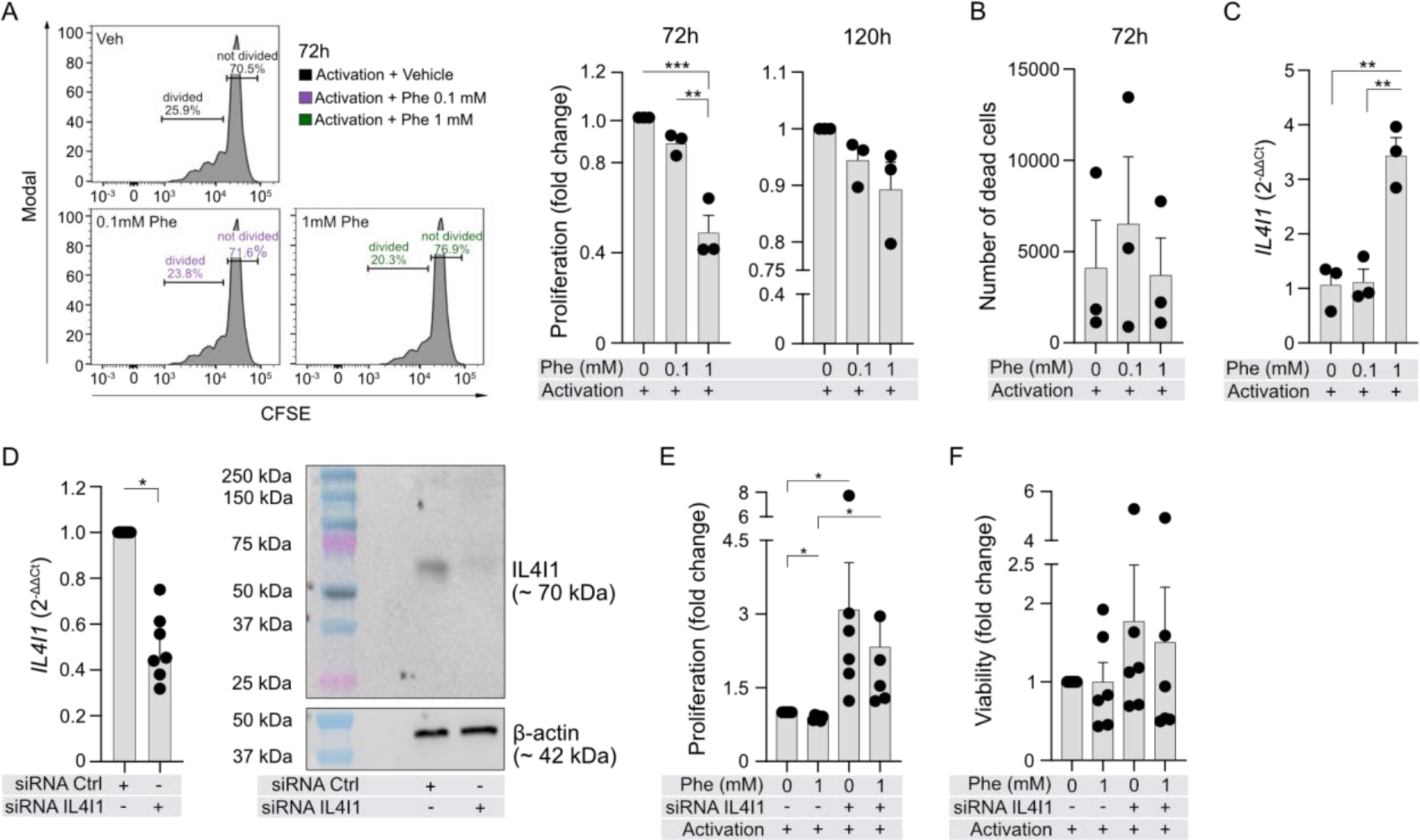
L-phenylalanine inhibits proliferation of human memory CD4^+^T cells by induction of Interleukin 4 induced 1 (IL4I1) enzyme. (**A**) Representative flow cytometry proliferation histograms (left) and cell proliferation quantification (right) of memory CD4^+^T cells incubated in full medium supplemented with Vehicle, 0.1mM or 1mM Phe and treated with CD2, CD3 and CD28 activation antibodies for 72h and 120h. (**B**) Number of dead memory CD4^+^T cells in the same experiments as in (A). (**C**) Expression of *IL4I1* mRNA in memory CD4^+^T cells following similar treatment as in (A). (**D**) *IL4I1* mRNA expression (left) and representative WB image of IL4I1 (right) in siRNA knockdown experiments in memory CD4^+^T cells. Cells were electroporated with *IL4I1* or negative Ctrl siRNA (300nM) and incubated for 24h. Data show 3 independent experiments in 6 different subjects. One outlier was identified using Grubbs’ test with α=0.05. (**E**) Proliferation of control siRNA (Ctrl) and IL4I1 siRNA treated memory CD4^+^T cells from 2 independent experiments in 5 different donors, incubated in full medium with/without 1mM additional Phe and treated with/without CD2, CD3 and CD28 activation antibodies for 24h before flow cytometry. Figure S2 shows non-activated conditions. (**F**) Viability of control siRNA (Ctrl) and IL4I1 siRNA treated memory CD4^+^T cells in the same experiments as in (E). (A-C) Data from three independent experiments in three different subjects. (A, B, E, F) Bar graph shows fold change as compared to activated Vehicle-treated cells. One-way ANOVA with Tukey correction was used in A and B; Unpaired t-test was used in C; Wilcoxon test was used in D-F. All data are presented as mean±SEM. *p<0.05, **p<0.01, ***p<0.001. Legend: Phe: L-phenylalanine.

In short, Phe in OXPHOS-impairing doses also limits proliferation of memory CD4^+^T cells via an IL4I1-dependent mechanism.

### L-Phenylalanine induces glycolysis and represses OXPHOS in activated human Th2, but not Treg cells analysed on a single cell level

Next, given that previous data revealed T cell-specific and systemic alterations in energy and amino acid metabolism in type 2 inflammatory diseases^38,45–48^, we studied the effect of Phe on the metabolism of subpopulations of human CD4^+^T cells, focusing on Th2 effector cells and Treg cells on the single cell level. First, to confirm that extracellular Phe enters the cells, we measured intracellular Phe in activated Th2 cells incubated in Phe-supplemented media, observing its significant increase **(Figure 4A)**. Next, we employed a recently developed method of assessing single cell energetic metabolism by profiling translation inhibition (SCENITH)^49^, following the protocol depicted in **Figure 4B** and panel provided in **Table S8**. Several of the analysed human CD4^+^T cell subsets, including Th2 (CD4^+^GATA3^+^FOXP3^-^ CCR4^+^) and Treg (CD127^-^CD25^++^FOXP3^+^) cells, highly increased the level of protein translation upon activation **(Figure 4C-4F)**, which corresponds with ATP production, as previously described.^49^ In Th2 cells, an increase in translation **(Figure 4F)**, glucose dependence **(Figure 4G)** and glycolytic capacity **(Figure 4H)** was observed upon activation, paired with a significant and substantial decrease in mitochondrial dependence **(Figure 4I)** as well as FAO and amino acid oxidation (AAO) capacity **(Figure 4J)**. Addition of Phe significantly increased glycolytic capacity of activated Th2 cells **(Figure 4H)** and profoundly decreased their mitochondrial dependence **(Figure 4I)**, which was reflected in a decrease of the general level of translation **(Figure 4F)**. In Treg cells, upon activation, we observed an increase in translation, which was slightly lower than in Th2 cells **(Figure 4E, 4F).** There was also an activation-induced increase in their glycolytic capacity **(Figure 4L)** and a reduction in mitochondrial dependence **(Figure 4M)**, but no changes in glucose dependence **(Figure 4K)** or FAO and AAO capacity **(Figure 4N)**. Interestingly, in Treg cells, Phe did not significantly affect any of the glycolytic or mitochondrial processes **(Figure 4K-4N)**.

**Figure 4.**
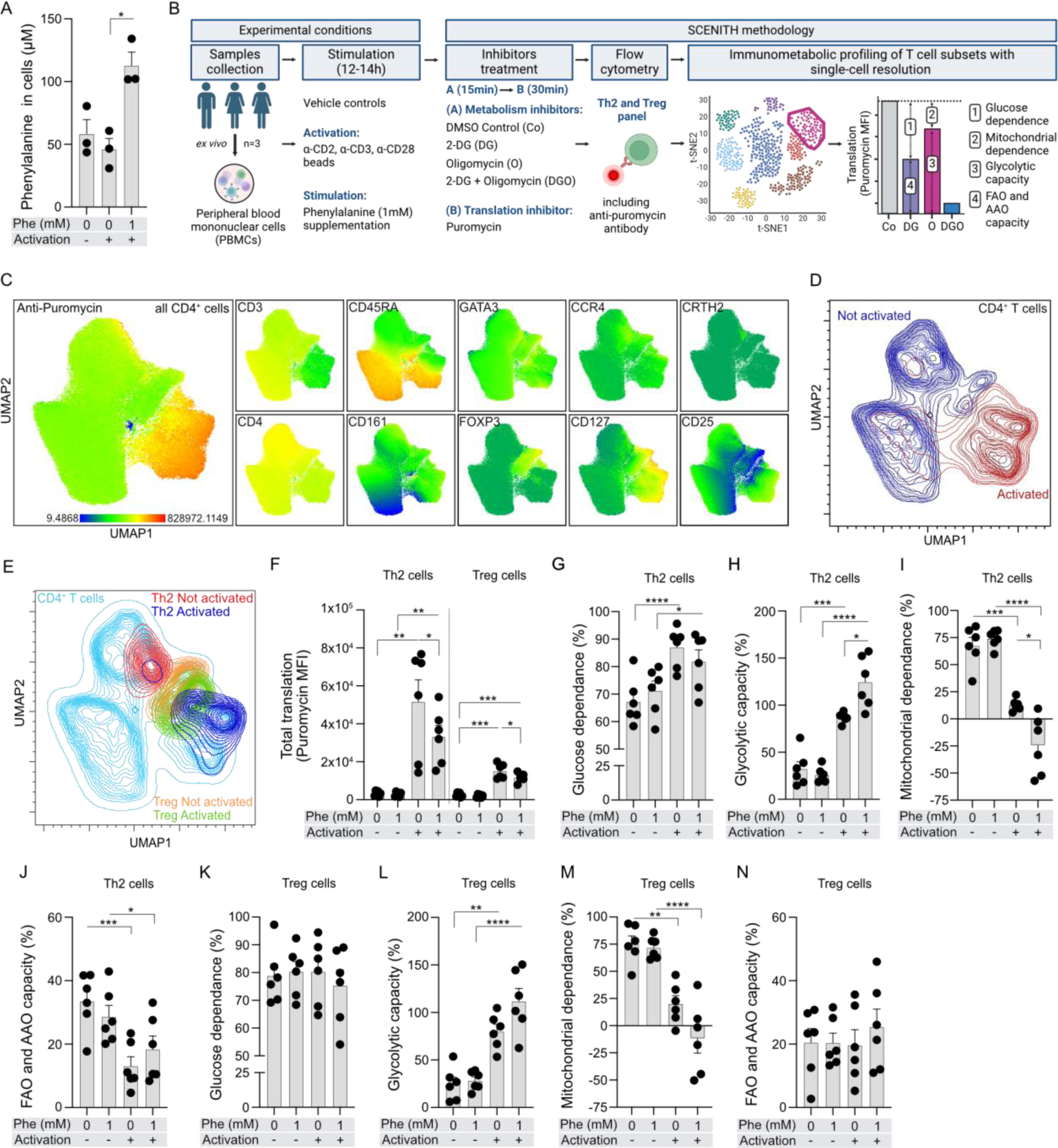
Single cell energy metabolism profiling revealing that L-phenylalanine increases glycolytic capacity and decreases mitochondrial dependence in activated human Th2 cells. (**A**) Phe uptake into Th2 cells. *In vitro* differentiated Th2 cells from 3 different donors were incubated in full medium with/without supplementation of 1mM Phe with/without CD2, CD3 and CD28 activation antibodies for 6h, and intracellular Phe was colorimetrically quantified in lysates. Data are analysed using paired t-test. (**B**) Schematic explaining workflow of Single Cell ENergetIc metabolism by profilIng Translation inHibition (SCENITH)^49^. PBMCs from three different healthy subjects were incubated in full medium with/without supplementation of 1mM Phe with/without CD2, CD3 and CD28 activation antibodies for 12-14h. Then, cells were treated with specified energy metabolism pathway inhibitors and stained with human Th2/Treg flow cytometry panel (Table S8) along with anti-puromycin antibody and measured by flow cytometry. Puromycin incorporation (MFI) in inhibitor treated samples enables glucose dependence, mitochondrial dependence, glycolytic capacity, and Fatty acid and Amino acid oxidation (FAO and AAO) capacity determination as per formulae provided in Arguello RJ *et. al.,* 2020. Figure created with Biorender.com. (**C**) Uniform Manifold Approximation and Projection (UMAP) plots of indicated protein expression in activated and non-activated CD4^+^T cells. FlowJo plugin UMAP (Ver 4.0.4) was used for analysis. (**D-E**) UMAP contour plots demonstrating activated and non-activated CD4^+^T (**D**) and Th2 and Treg (**E**) cells. (**F**) Total translation in Th2 and Treg cells. One-way ANOVA with Fishers LSD test was used for analysis. (**G-N**) Glucose dependence (**G, K**), glycolytic capacity (**H, L**), mitochondrial dependence (**I, M**), and FAO and AAO capacity (**J, N**) in Th2 and Treg cells, respectively. (F-N) Each dot represents one technical replicate per donor (n=3 donors, 2 replicates each). (G-N) For comparing dependencies and capacities, One-way ANOVA with Sidaks’ multiple comparison correction was used. Bars represent mean±SEM. *p<0.05, **p<0.01, ***p<0.001, ****p<0.0001. Legend: Phe: L-phenylalanine.

These data suggest that high intracellular levels of Phe might repress the global ability of Th2 cells to produce ATP, essential for translation. In addition, Phe causes profound metabolic reprogramming stimulating glycolysis and blocking mitochondrial OXPHOS, especially in activated CD4^+^GATA3^+^CCR4^+^FOXP3^-^Th2 cells.

### L-Phenylalanine inhibits Th2 cell proliferation, transcription factors, cytokines and activation markers

Following the effect of Phe on energy metabolism and proliferation of human memory CD4^+^T cells, as well on Th2 metabolism on a single cell level, we next studied in detail the effect of Phe on metabolism, proliferation, phenotype, and function of *in vitro* highly differentiated human Th2 cells^50^ **(Figure S3)**, comparing it to Arg. Similar to Phe-treated, activated total and memory CD4^+^T cells **(Figure 2)** and Th2 cells **(Figure 4H)**, here, we observed increased levels of inducible glycoATP in activated *in vitro* differentiated Th2 cells incubated in 1mM of additional Phe **(Figure S4A)**. However, we did not note any major influence of Phe on mitoATP levels **(Figure S4B)**. In contrast, additional Arg did not have any effect on *in vitro* differentiated Th2 cell energy metabolism **(Figure S4C, S4D)**. Interestingly, supplementation with 1mM of additional Phe significantly reduced proliferation of activated Th2 cells **(Figure 5A)** while increasing their viability **(Figure 5B)**. In non-activated conditions, Phe increased proliferation of Th2 cells **(Figure S5A)**, with no effect on their viability **(Figure S5B)**. Next, we tested the influence of Phe and Arg on expression of major transcription factors and their co-factors, type 2 cytokines, surface markers, and other molecules responsible for Th2 differentiation and activation **(Figure 5C-5E, Figures S6-S8)**. We observed significantly reduced mRNA expression of type 2 transcription factors^51–53^ such as *MTOR*, *RICTOR* and *RAPTOR*, *BATF* and *BACH2* **(Figure 5C)**, cytokines *IL4*, *IL5*, *IL13* **(Figure 5D)** and surface markers such as *CD69 and PTGDR2* (CRTH2) **(Figure 5E)** in activated Th2 cells incubated with increasing concentrations of Phe. Phe did not affect mRNA expression of *GATA3*, *STAT6*, *IFNG*, *IL4I1*, *IL2RA*, or *PDCD1* **(Figure 5C-5E)**. These effects of Phe were not observed in non-activated Th2 cells **(Figure S6)**. Importantly, Arg did not affect the mRNA expression of these molecules in Th2 cells at the same doses **(Figures S7, S8)**. Finally, since it has been previously reported that CD161 is an important marker associated with pathogenicity of Th2 cells (so-called Th2a cells) in humans^54^, we used flow cytometry to assess whether its expression is influenced by Phe. Indeed, we observed a significant reduction of CD3^+^CD4^+^CCR4^+^GATA3^+^CD161^+^ Th2 cell frequency in activated conditions upon treatment with Phe **(Figure 5F, Figure S9).**

**Figure 5.**
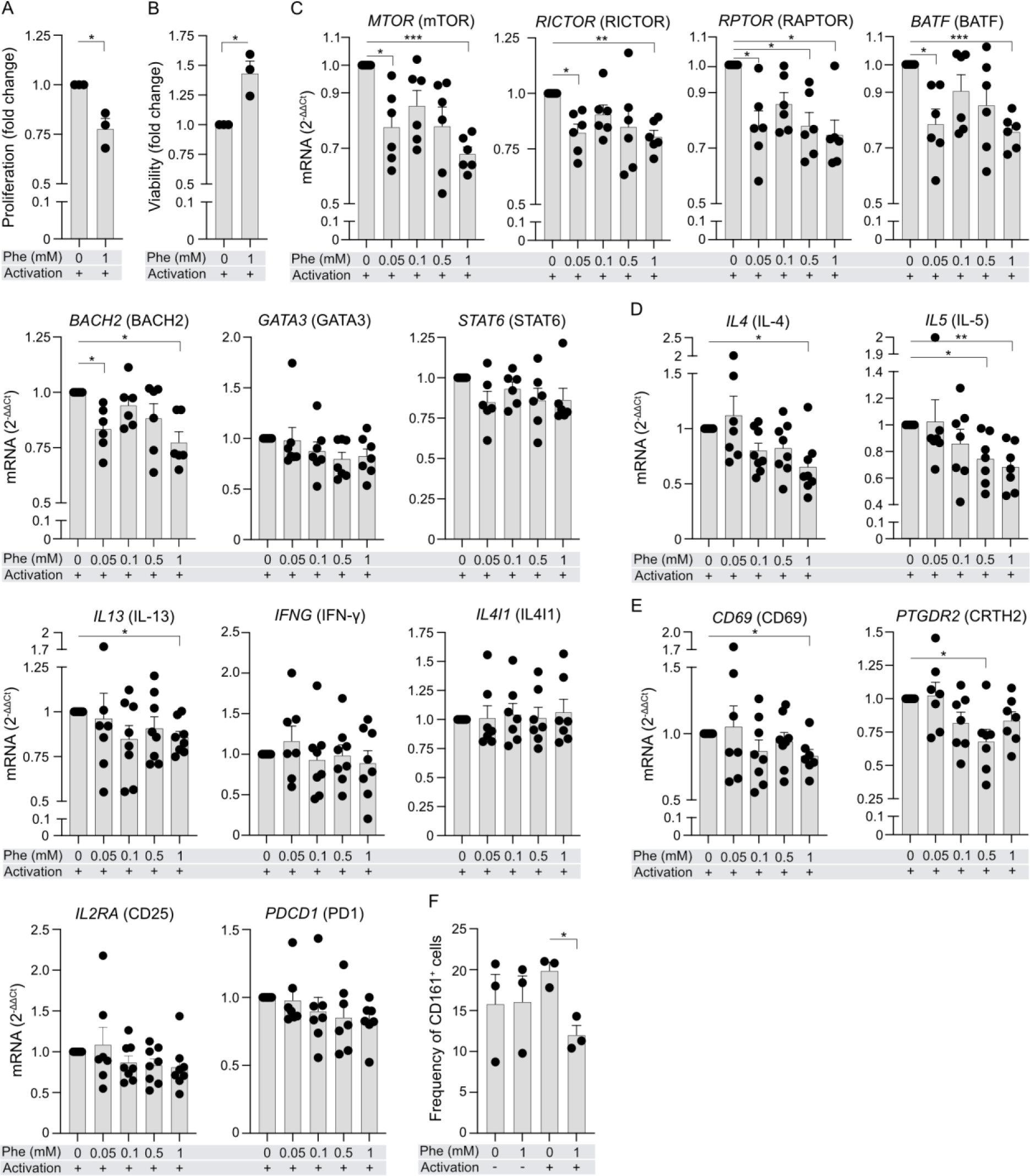
L-phenylalanine inhibits Th2 cell proliferation and expression of type 2 transcription factors, cytokines, activation, and pathogenicity markers. (**A-B**) Proliferation (**A**) and viability (**B**) of *in vitro* differentiated Th2 cells subjected to high doses of Phe. Th2 cells were incubated in full medium with/without additional supplementation of 1mM Phe with/without CD2, CD3 and CD28 activation antibodies for 24h. Both parameters were assessed by flow cytometry. Bar graphs show fold changes compared to Vehicle-treated, activated cells. Paired t-test was used for analysis. n=3 different donors. Figure S5 depicts non-activated conditions. (**C-E**) mRNA expression of transcription factors (**C**), cytokines and enzymes (**D**), and surface markers (**E**) in *in vitro* differentiated Th2 cells treated with increasing doses of Phe. Th2 cells were incubated in full medium supplemented with Vehicle, 0.05mM, 0.1mM, 0.5mM and 1mM Phe with/without CD2, CD3 and CD28 activation antibodies for 24h. Following incubation, mRNA expression was determined using qRT-PCR. n=6-8 different donors. Figure S6 shows non-activated conditions. (**F**) Frequency of CD3^+^CD4^+^CCR4^+^GATA3^+^CD161^+^Th2 cells following incubation with/without supplementation of 1mM Phe with/without CD2, CD3 and CD28 activation antibodies for 24h and staining with panel in Table S8 for flow cytometric assessment. (n=3 different donors). (C-F) Data are analysed using One-way ANOVA with Dunnett’s correction. Each dot represents one donor. Bars represent mean±SEM. *p<0.05, **p<0.01, ***p<0.001. Legend: Phe: L-phenylalanine.

In summary, in *in vitro* differentiated human Th2 cells, Phe, but not Arg, affects TCR activation-induced glycolysis, proliferation and transcription of several key molecules involved in differentiation and function of Th2 cells, including markers of their pathogenicity.

### Low intracellular levels of L-Phenylalanine in human memory CD4^+^T effector cells characterize severe allergic patients with systemic type 2 inflammation

Metabolic dysregulation of T cell subsets has been observed in several acute and chronic inflammatory diseases^55–57^; however, for type 2 inflammation and disease in humans, it has not been thoroughly studied.^48,57^ We and others demonstrated that patients with severe forms of allergic diseases, such as severe allergic asthma or severe allergic rhinitis, exhibit metabolic differences in serum and cellular secretomes in comparison to healthy subjects and patients with mild allergies.^47,58^ In addition, circulating and tissue residing memory Teff cells and Treg cells display a wide range of abnormalities in function and phenotype in allergic diseases.^59^ Finally, having observed here a strong influence of Phe on Th2 energy metabolism, cell proliferation and phenotype, we hypothesized that the metabolism of circulating memory Teff and Treg cells in allergy might reflect, to some extent, their changed functionality, systemic inflammatory milieu, or clinical characteristics.

We recruited non-allergic, mild, and severe allergic individuals to olive pollen out of the pollen season (Cohort A) **(Tables S9, S10)**. Cohort A consisted of clinically well described subjects at different stages of allergic sensitization and symptoms.^60,61^ We first characterized their immunological status in great detail by analysing CD4^+^T cell and ILC compartments, and serum proteomics to assess the level of cellular and systemic type 2 inflammation. Severe allergic patients had higher number of pathogenic memory CRTH2^+^CD161^+^Th2a cells^31,54^ compared to non-allergic controls and mild allergic patients **(Figure 6A)**, although, their total Th2 and Treg cells were not significantly different **(Figure S10A)**. Interestingly, they also had higher number of CRTH2^+^Treg and PD1^+^Treg cells **(Figure 6B)** which have been reported by us and others to have inefficient regulatory capacity.^62–64^ In addition, severe allergic patients had high numbers of ILC2 and ILC3 **(Figure 6C, Figure S10B)**, in agreement with previous reports in other allergic cohorts.^65–67^ In line with that, by performing targeted proteomics, we found several proteins to be differentially expressed in serum of severe allergic patients compared to non-allergic and/or mild allergic subjects **(Figure 6D, Figure S11)**. IL-5, and proteins involved in cell proliferation namely BOC and PAPPA were significantly elevated in severe allergic patients **(Figure 6D)**. These patients also had lower concentration of Th1-related cytokines such as IL-12, CSF-1 or GZMH and Treg-associated hepatocyte growth factor (HGF) **(Figure S11)**.

**Figure 6.**
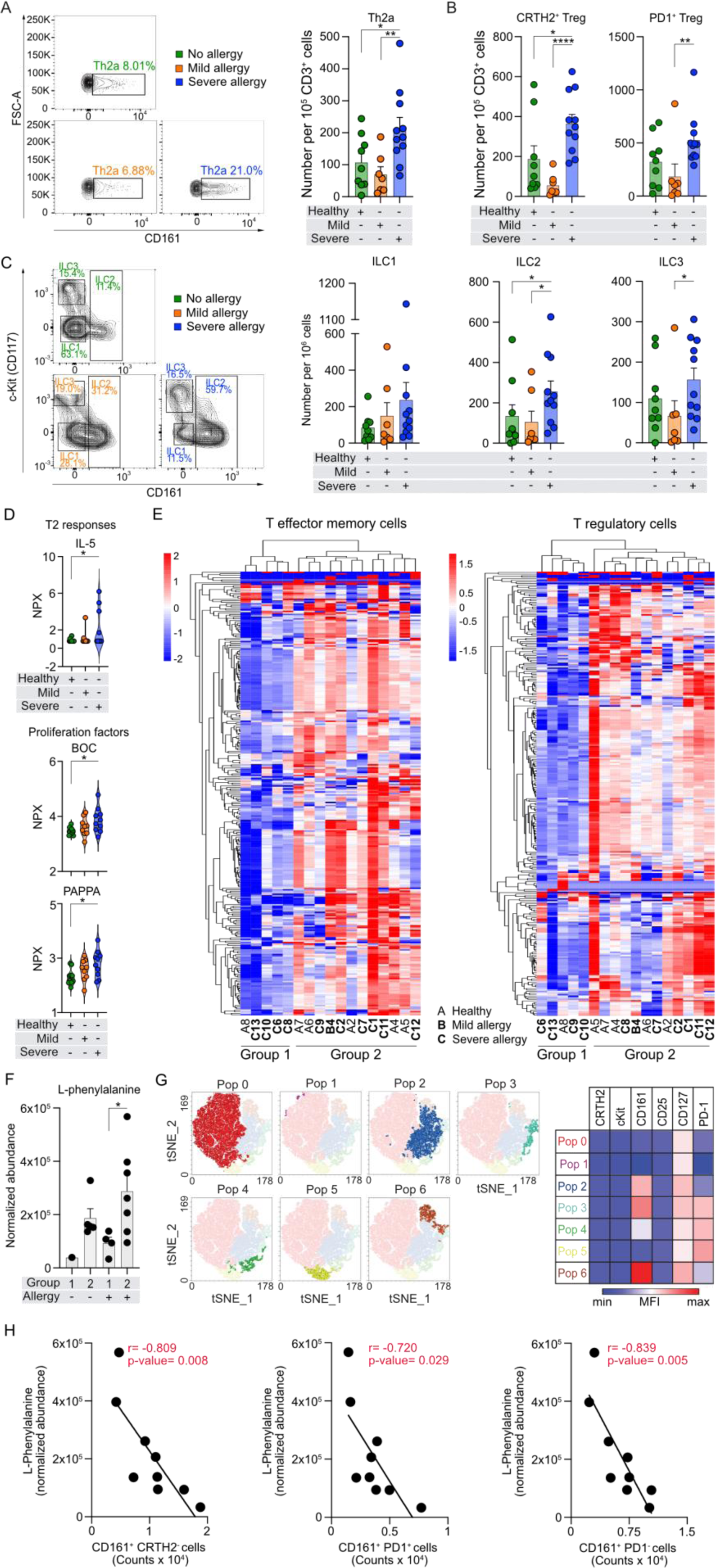
Low intracellular L-phenylalanine levels in pathogenic memory CD4^+^T effector cell populations in severe allergic patients. (**A-C**) Representative flow cytometry dot plots of Th2a cells (**A**-left) and ILC1, ILC2, and ILC3 (**C**-left). Number of Th2a cells (**A**-right), memory CRTH2^+^Tregs and PD1^+^Tregs (**B**) and ILC1, ILC2, and ILC3 (**C**-right) in controls (n=9), patients with mild (n=7) and severe (n=11) allergy (Cohort A). (**D**) Differentially expressed proteins in serum of controls (n=10), mild (n=9) and severe (n=10) allergic patients (Cohort A) assessed with PEA technology and presented as NPX. (**E**) Heatmaps with hierarchical clustering analysis of all metabolites measured in memory CD4^+^Teff (n=195, left) and Treg (n=233, right) cells in controls (n=6) and allergic (n=11) subjects. A-controls, B-mild allergy, C-severe allergy (from Cohort A). (**F**) Normalized abundance of Phe in Teff cells in group 1 (control, n=1; severe allergy, n=4) and 2 (control, n=5; mild allergy, n=1, severe allergy, n=6) (from Cohort A). (**G**) tSNE plot of unbiased 2-dimensional flow cytometric analysis of memory CD4^+^Teff cells (CD3^+^CD4^+^CD45RA^-^CD127^+^CD25^-^) from patients with severe allergy (subset of cohort A, n=9) identifying six subpopulations based on CRTH2, cKit, CD161, and PD-1. (**H**) Pearson correlation of normalized abundance of intracellular Phe, in memory CD4^+^Teff cells from patients with severe allergy (n=9), with counts of CD161^+^ populations within memory CD4^+^Teff cells (from Cohort A). Mann-Whitney U test (A-C), One-way ANOVA with Fishers LSD test (D) and Unpaired t-test (F) were used to compare differences among groups. Graphs represent mean±SEM. *p<0.05, **p<0.01, ***p<0.001. Clinical characteristics of Cohort A are shown in Tables S9 and S10. NPX: Normalized Protein eXpression; Pop: population; Phe: L-phenylalanine.

Next, we sorted the circulating memory CD4^+^Teff and Tregs from a subset of Cohort A patients (n=11) and compared them to non-atopic, non-allergic healthy controls (n=6), and analysed their metabolomes. We did not find any significant differences between healthy and allergic patients either in Teff or in Treg cells metabolomes **(Tables S11, S12)**. However, we identified two groups of subjects, whose samples clustered together **(Figure 6E)**. Most metabolites in group 1, consisting of 80% of severe allergy patients, were less abundant than in group 2, consisting of 50% of severe patients. These findings suggest that there are 2 distinctive cellular metabolic phenotypes in both cell subsets. Interestingly, group 1 and 2 consisted of mostly the same subjects in both cell types potentially suggesting their metabolic profiles are linked. Importantly, among them, Teff cells from allergic patients in group 1 had significantly lower levels of Phe **(Figure 6F)**, Arg and Leu/Ile **(Figure S12)**. To gain a clearer understanding of the differences behind this grouping, especially considering that memory CD4^+^Teff cells are a heterogeneous group, we analysed their phenotype in a multidimensional and unbiased way. We identified seven and eight different populations within CD4^+^CD127^++^CD25^-^Teff cells in allergic patients **(Figure 6G)** and healthy controls **(Figure S13A, S13B)**, respectively, mostly differing in expression of CD161 and PD-1. Next, we correlated the abundance of amino acids with the identified subpopulations. Notably, in allergic patients, we found strong negative correlations between levels of intracellular Phe and counts of CD161^+^ populations **(Figure 6H)**, but not CD161^-^ populations **(Figure S13C)**, which was not the case in healthy controls **(Figure S13D)**.

In summary, memory CD4^+^Teff cells in a subset of patients with a clinically diagnosed severe form of allergy contain lower amounts of Phe. Additionally, we noted lower level of intracellular Phe and higher expression of CD161, a Th2a pathogenicity marker, on the surface of memory CD4^+^Teff cells in patients with severe allergy.

### Extracellular L-phenylalanine decreases expression of large-neutral amino acid transporters in Th2 cells *in vitro,* which is reflected *in vivo* in allergic patients

Since we observed an influence of intracellular and extracellular Phe on metabolism, phenotype, proliferation, and function of human CD4^+^T cells, memory CD4^+^T cells, and Th2 cells, and we found that lower intracellular level of Phe in Teff cells correlates negatively with phenotypes and frequencies of Th2 cells subpopulations in allergic patients, we next explored expression of main molecules involved in Phe metabolism and transport in other cohorts of patients with allergic diseases.

First, we analysed Phe metabolism and transport in previously published RNA-Seq data from Th2 cells (CD3^+^CD4^+^CD45RA^-^CD25^-^CCR4^+^) sorted from patients with allergic asthma (n=37) and allergic rhinitis (n=25) (allergic to respiratory and other allergens) or healthy controls (n=15) (Cohort B).^68^ Interestingly, we found that several of the top significant Gene Ontology (GO) processes enriched in Th2 cells were related with metabolic processes in patients with allergic asthma **(Figure 7A, Table S13)** and allergic rhinitis **(Figure S14A, Table S14)**. In detail, there was an upregulation of amino acid metabolism **(Figure 7B, S14B, Tables S15-S16)** and downregulation of L-threonine and L-arginine pathways and transport was observed in allergic asthma **(Figure 7C, Table S17)** and allergic rhinitis patients **(Figure S14C, Table S18)**, respectively. Several genes responsible for phenylalanine metabolism and transport were included in these pathways **(Table S19)** and were significantly changed in Th2 cells from asthma patients **(Figure 7D-7E)**. We noted that *SLC7A5* (LAT1), *SLC7A8* (LAT2), and *SLC3A2* (CD98), encoding for the additional part of LAT1 and LAT2 heterodimer complexes, were all significantly downregulated in patients with allergic asthma as compared to healthy controls **(Figure 7D-7E)**, suggesting decreased transport of Phe into the Th2 cells in allergy. In addition, we noted that *GOT1* and *GOT2*, involved in biosynthesis of phenylpyruvate from Phe, were significantly upregulated in allergic asthma patients in Th2 cells **(Figure 7D-7E)**. Also, other genes encoding for enzymes indirectly involved in metabolism of Phe, such as pterin-4 alpha-carbinolamine dehydratase 2 (PCBD2), catechol-O-methyltransferase (COMT) and kynurenine aminotransferase 1 (KYAT1), were significantly upregulated, altogether suggesting potentially faster turnover of intracellular Phe in Th2 cells of patients with allergic asthma. Although we observed a trend of reduced expression of these genes in a similar assessment of allergic rhinitis patients against healthy controls, this decrease was not significant (**Figure S14D**). We also analysed gene expression of molecules belonging to Phe metabolism and amino acid transport in total CD3^+^T cells in our own previously reported cohort (Cohort C)^46^ consisting of healthy non-allergic controls (n=8), and patients with mild (n=9) and severe (n=7) allergic asthma (GEO: GSE224253). We observed significantly reduced expression of multiple key genes involved in Phe metabolism in total CD3^+^T cells from severe allergy patients in comparison to controls **(Figure S14E).** With the exception of *SLC7A5*, we did not observe significant reduction in the expression of Phe metabolism genes in CD3^+^T cells from patients with mild allergy in comparison to controls **(Figure S14E, S14F)**.

**Figure 7.**
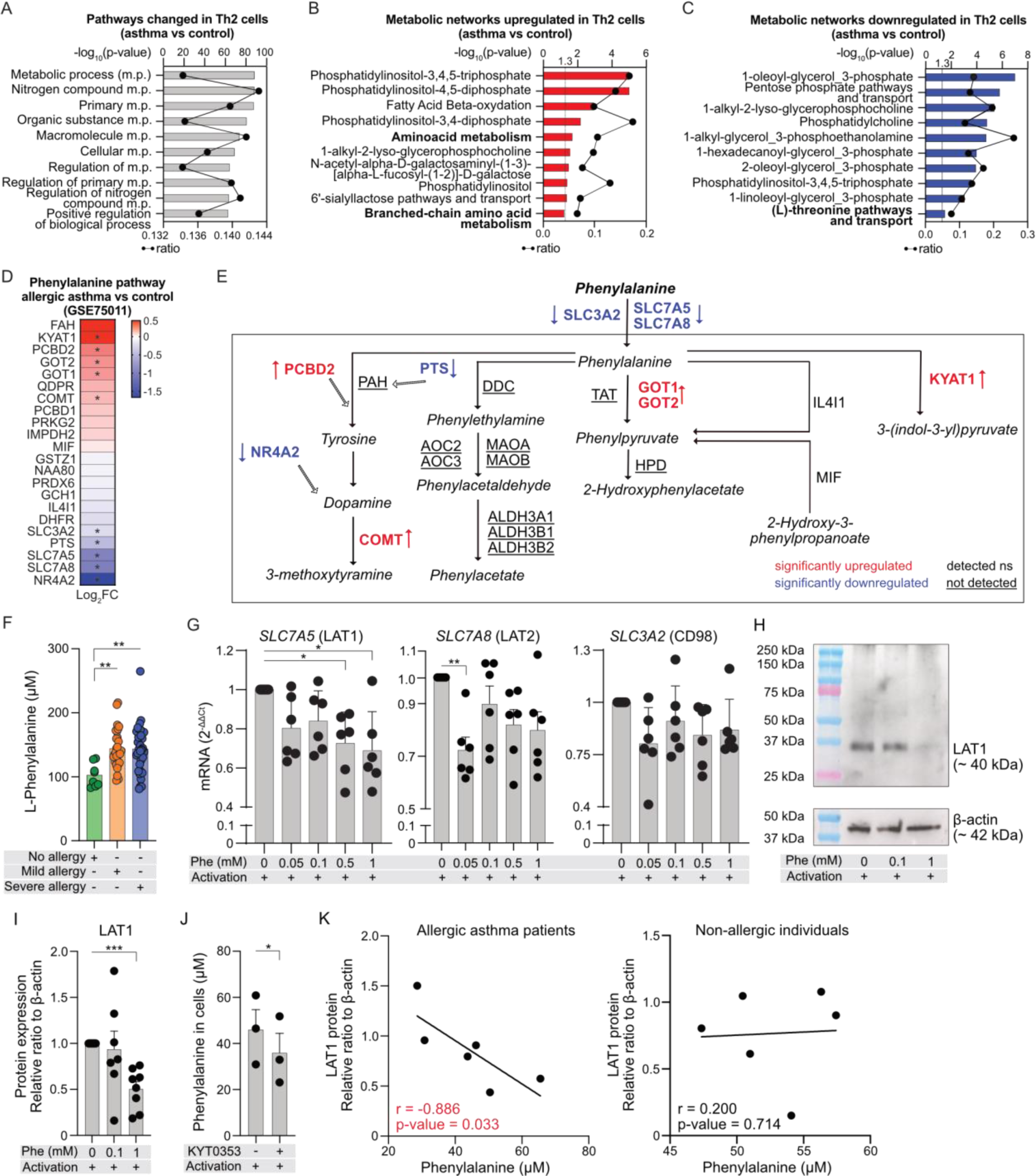
Decreased expression of large amino acid transporters (LAT) in Th2 cells of allergic patients correlates with elevated serum levels of L-phenylalanine. (**A-C**) Top significantly enriched pathways (**A**), significantly upregulated (**B**) and significantly downregulated (**C**) metabolic networks within differentially expressed genes (DEG, p<0.05) in allergic asthma patients compared to controls (control n=15, allergic asthma n=37) from GSE75011 (Cohort B)^68^. Black line represents ratio of genes in experiment over complete pathway set. All data available in tables S13, S15 and S17, respectively. (**D**) Phe metabolism and transport pathway heatmap showing fold change (Log_2_FC) of DEGs in allergic asthma patients (n=37) compared to controls (n=15) from GSE75011 (Cohort B)^68^. *p<0.05. Pathway curated and adapted from GSEA and MSigDB Database (Table S19). (**E**) Phe metabolism and transport schematic highlighting DEGs in Th2 cells of allergic asthma and controls (GSE75011) (Cohort B)^68^. Significantly upregulated (red) and downregulated (blue) genes; detected but not significantly different (black); not detected in original dataset (black and underlined). Adapted from KEGG pathway. (**F**) Serum Phe concentration in controls (n=8), mild (n=30) and severe (n=37) allergic patients (Cohort D) quantified by targeted metabolomics^101^. Kruskal-Wallis test was used for analysis. (**G**) *SLC7A5* (LAT1)*, SLC7A8* (LAT2) and *SLC3A2* (CD98; LAT3) mRNA expression in *in vitro* differentiated Th2 cells treated with additional Phe with/without CD2, CD3 and CD28 activation for 24h. Following incubation, mRNA expression was determined using qRT-PCR (n=6-8 different donors). (**H-I**) Representative WB image of LAT1 expression (**H**) and LAT1 protein quantification in 8 different donors (**I**) in *in vitro* differentiated Th2 cells treated with additional Phe with/without CD2, CD3 and CD28 activation for 24h. Expression of LAT1 presented as relative ratio normalized to β-actin. (**J**) Phe uptake into *in vitro* differentiated Th2 cells was quantified colorimetrically. Cells were incubated in full medium with/without SLC7A5 inhibitor (KYT0353) and activation of CD2, CD3 and CD28 for 6h. Data were analysed using paired t-test (n=3 different donors). (**K**) Spearman correlation of serum Phe concentration and relative LAT1 expression in CD4^+^T cells in allergic asthma patients (left) and controls (right) (Cohort E). (G and I) Data are analysed by One-way ANOVA with Dunnett’s correction. (F-J) Bars represent mean±SEM. Each dot represents one donor. *p<0.05, **p<0.01, ***p<0.001. Cohorts D and E are characterized in Tables S20 and S21, respectively. Legend: Phe: L-phenylalanine.

Next, to determine if serum levels of Phe were altered, we measured Phe concentration in a new, large cohort of subjects including healthy controls (n=8), mild (n=30) and severe (n=37) allergic patients enrolled in the same hospital and with the same clinical criteria as those from the Cohort A (Cohort D, **Table S20**). We observed that mild and severe allergic patients had higher levels of Phe in serum compared to non-allergic individuals **(Figure 7F)**. To understand the relationship between the lower abundance of intracellular Phe in a subset of patients with severe allergy, lower expression of genes encoding for Phe transporters in RNA-Seq data of Th2 cells in allergic asthma, and higher concentration of Phe in the serum of severe allergic patients, we analysed an influence of extracellular Phe on gene expression of all parts of the LAT complexes. We found that increasing concentrations of extracellular Phe, but not Arg, significantly decreases mRNA expression of *SLC7A5* and *SLC7A8*, but not *SLC3A2* in *in vitro* cultured Th2 cells **(Figure 7G, S15)**. We confirmed this finding on the protein level, demonstrating decreasing expression of LAT1 with increasing doses of Phe in the culture medium **(Figure 7H-7I)**. In addition, we confirmed that LAT1 is needed for the transport of Phe into the activated Th2 cells, since LAT1 inhibitor KYT0353 decreased uptake of Phe **(Figure 7J)**. Finally, we measured LAT1 expression in CD4^+^T cells and Phe levels in serum from patients with allergic asthma and healthy controls from our previous study^69^ (Cohort E) **(Table S21)**. We found a negative correlation between serum Phe and LAT1 expression only in allergic asthma patients **(Figure 7K)**.

In summary, these data suggest that serum Phe levels in allergic patients are significantly elevated, and Phe dose-dependently decreases the expression of its main transporter LAT1 in Th2 cells. This phenomenon, together with increased expression of enzymes metabolizing Phe in Th2 cells, might further translate to lower intracellular Phe levels in Th2 cells and their expansion and pathogenicity in allergic diseases.

## Discussion

In T cells, metabolic checkpoints act as critical molecular switches that ensure an efficient transition of metabolism from one state to another, precisely reflecting their developmental needs and function.^70^ In our study, by using *ex vivo* metabolomics, lipidomics, single cell metabolism profiling and serum proteomics, combined with *in vitro* molecular and functional approaches, we have shown that Phe can serve as a checkpoint in human memory CD4^+^T cells and Th2 cells due to its pleiotropic effects on activation, energy metabolism, proliferation, phenotype, and function. We have also demonstrated that in humans with severe allergic diseases this checkpoint is impaired and the pro-inflammatory phenotype of pathogenic Th2 cell subsets may be facilitated by a decreased abundance of intracellular Phe, resulting from its impaired transport, and accelerated intracellular processing.

Metabolic specialisation of human and mouse Teff subsets and Treg upon activation have been continuously demonstrated, showing higher dependence on glycolysis in Teff subsets and on FAO and OXPHOS in Tregs^6,9,71^, although more and more studies demonstrate that this metabolic reprogramming is tissue-, nutrient- and context-dependent.^10,11^ In fact, T cells show metabolic plasticity depending on the environment and accessibility of nutrients. *In vivo*, when glucose level is much lower than in *in vitro* experiments, their energy metabolism, proliferative capacity, and effector response depend on other nutrients, such as amino acids or fatty acids.^12,20,72^ It was elegantly demonstrated among others by combination of *in vivo* stable isotope infusion and tracing with bioenergetic profiling.^73^ Such experiments, however, have limited use in humans. Therefore, employing combination of *ex vivo* cell sorting with high resolution metabolomics, as well as flow cytometry-based single cell energy metabolism profiling methods have a potential to explore metabolism of human T cells. Hence, we first utilised *ex vivo* cell sorting and metabolomics to study circulating human memory CD4^+^T cell compartment, focusing on previously unexplored comparison between human memory CD4^+^CD45RA^-^CD127^++^CD25^-^Teff and CD4^+^CD45RA^-^CD127^-^CD25^+^Treg cells. Our finding that metabolomes of these cells at the steady state differ substantially is quite striking, potentially reflecting the differences in their phenotype and function. This adds to the previous findings showing that metabolome of *ex vivo* purified human naïve CD4^+^T cells differs from metabolome of naturally occurring CD4^+^CD25^+^Treg^10^ or thymus-derived Treg cells.^74^ We found enrichment in various amino acids from different metabolic groups in both cell types, seemingly taking part not only in protein synthesis and degradation, but also in glycolysis, gluconeogenesis, pentose phosphate, fatty acid, and lipid metabolism pathways. In each cell type, however, particular amino acids might be used by different metabolic pathway^19^, considering their significantly different abundance. This is in line with previous observations of differential enrichment in amino acids in Th1, Th17 and T follicular helper (Tfh) cells.^75^

Likewise, we observed a reduction in histidine, proline and Arg in CD4^+^Teff compared to Tregs, suggesting their faster utilisation as it was demonstrated in activated human CD4^+^T cells.^20^ We also confirmed in the single cell energy profiling experiments that *ex vivo* activated Teff cells, particularly Th2 cells, tended to have higher translation-used ATP production than Treg cells, suggesting differential anabolic and catabolic usage of available amino acids in both cell types.

Amino acids guide cellular functions through their selective sensing and acquisition which control many biosynthetic processes and steer energy metabolism.^19^ Here, we showed that supplementation of Arg during activation increases glycolysis and maximum respiratory capacity in freshly sorted human CD4^+^T cells but does not change glycoATP or mitoATP production in *in vitro* differentiated human Th2 cells. Arg is transported in T cells via cationic amino-acid transporter CAT-1 (SLC7A1).^20,76^ Interestingly, in naïve human CD4^+^T cells upon activation, in the conditions of very high extracellular Arg supply (4mM), Arg is quickly metabolized, feeding gluconeogenesis, limiting glucose uptake, and increasing OXPHOS.^20^ This in turn, limits differentiation, but increases survival and maintains them in the memory like state.^20^ Our data suggests that in a situation of lower Arg levels (around 1mM), it can still feed into glycolysis and OXPHOS, at the early differentiation stage of CD4^+^T cells, but not after terminal differentiation to Th2 cells. In contrast, Phe, transported via LAT1 or LAT2^77,78^, in the same concentrations increased activation-induced glycolysis in total and memory human CD4^+^T cells, glycoATP production in *in vitro* differentiated Th2 cells and glycolytic capacity in *ex vivo* treated Th2 cells. It underlines Phe-induced glycolysis importance at different stages of CD4^+^T cells differentiation. In highest concentrations, however, in total and memory CD4^+^T cells Phe blocked activation-induced OXPHOS, did not activate mitoATP production in *in vitro* differentiated Th2 cells and downregulated mitochondrial dependence in *ex vivo* treated Th2 cells. This observation is interesting, especially since we noted that the same concentrations of Phe limited memory CD4^+^T cells and Th2 cells proliferation and type 2 cytokine production, while increasing their viability. It all suggests that during recall responses Phe might regulate the human memory Th2 cells by shifting its metabolites between OXPHOS and glycolysis.

In human T cells, phenylalanine hydroxylase (PAH), the main enzyme in Phe metabolism, is not expressed.^68,79^ We found, however, that they express other enzymes metabolizing Phe. PCBD2 catalyzes conversion of Phe to tyrosine.^80^ We found tyrosine in metabolomes of Teff and in Treg cells, and upregulation of PCBD2 gene in Th2 cells in allergic patients, suggesting that this pathway might be active. GOT1 and GOT2^81^, as well as IL4I1^82^ can catalyze conversion of Phe to phenylpyruvate. We also found phenylpyruvate in the memory Teff and Treg cells, as well as upregulated expression of GOT1 and GOT2 in Th2 cells of allergic asthma patients, suggesting that they might be involved in faster turnover of Phe driving pathogenic Th2 cells activity. IL4I1 can catalyse the conversion of Phe to phenylpyruvate liberating H_2_O_2_ and ammonia^43^, as well as can convert tryptophan to indoles and kynurenic acid activating aryl hydrocarbon receptor (AHR).^28^ In CD8^+^T and CD4^+^T cells, external IL4I1 or intracellular upregulation of IL4I1 can limit their proliferation via temporary downregulation of TCR complex.^28,43,44^ CD4^+^T cells, along with inhibition of proliferation^43^, external IL4I1 also facilitates the differentiation of naïve CD4^+^T cells into Treg cells.^27^ Here, we found that high levels of Phe increased expression of *IL4I1* in memory CD4^+^T cells and, in agreement with previous findings in Th17 cells^44^, that genetic inhibition of IL4I1 increased proliferation of memory CD4^+^T cells.

Amino-acid-induced Raptor-mTORC1 activity is required in CD4^+^T cells to regulate TCR-induced glycolysis, lipid biosynthesis and OXPHOS, needed to exit from quiescence and initiate differentiation into various T helper subsets, including Th2 cells.^35^ Lack of Raptor decreases IL-4 production by Th2 cells *in vitro* and diminishes allergic airway inflammation *in vivo*.^35^ Also, Rictor-mTORC2 pathway is important for Th2 cells reprogramming.^23,36^ Amino acids also regulate Treg function via licensing TCR-induced mTORC1 activation.^83^ Accordingly, we noted an overrepresentation of mTOR pathway in metabolomes of human *ex vivo* memory CD4^+^Teff and in Treg cells, paired with the enrichment of many amino acid pathways. In addition, we found that increased concentrations of Phe, but not Arg, upon TCR activation decreased expression of *MTOR*, *RPTOR* and *RICTOR* under established Th2 conditions, which was coupled with decreases in expression of all type 2 cytokines, *IL4*, *IL5* and *IL13*, surface activation marker *CD69* and Th2-pathogenicity markers such as *PTGDR2* (CRTH2) and CD161 protein. Since these phenomena were coupled with an induction of IL4I1 pathway by Phe, they might result from the negative impact of IL4I1 on mTORC1 signalling^27^, potentially via activation of AHR.^28,44^ Interestingly, high doses of Phe also increased glycolysis and survival in Th2 cells *in vitro*, as well as increased glycolytic capacity, but decreased OXPHOS and translation in *ex vivo* studied human Th2 cells. It might all suggest that high concentrations of Phe present during TCR activation of Th2 cells might reduce mTORC1 signalling either via IL4I1 or reduced OXPHOS, leading to decreased proliferation, transcription, and translation of type 2 cytokines.

FA synthesis, oxidation, and signalling are also very important in Th2 cell biology.^37,84^ We noted a general enrichment in glycerophospholipid metabolism in memory Teff, as well as fatty acid biosynthesis in Treg cells metabolomes. There was also an upregulation of phosphatidylinositol and FAO pathways in transcriptome of circulating human Th2 cells. It suggests that longevity and/or function of human circulating memory CD4^+^ and Th2 cells might depend on fatty acid uptake and metabolism, as it is in the case of tissue resident Th2 cells.^45,85^ Various subsets of Th2 cells at the steady state or during type 2 inflammation express transcription factors, enzymes and receptors involved in lipid signalling and metabolism, including CRTH2.^33,85–87^ They are all involved to a different extent in production of type 2 cytokines and cell trafficking. Here, we show that TCR-activation in human *ex vivo* studied Th2 cells, but not in Treg cells decreases FAO, which coincides with a decrease in expression of CRTH2 protein and transcript (*PTGDR2)*. Importantly, in TCR activated Th2 cells amino acid and fatty acid metabolism are strongly connected by mTORC1.^88^ Our severe allergic patients had significantly more CRTH2-expressing Th2a cells, Treg cells and ILC2 cells, as noted before.^54,62^ This upregulation of pathogenic CD4^+^T and ILC2 populations in severe, but not mild patients could potentially be attributed to a long term persistence of several CD4^+^ memory T cell clones specific to several epitopes and different allergens, or more severe inflammation at the mucosal sites, not fully resolved even in the absence of the allergen.^89^ Therefore, our findings that excessive Phe inhibits pathogenic Th2 cells might suggest its additional effect on fatty acid metabolism via mTOR, which is impaired in allergy.

The increase in amino acid demand following T cell activation is satisfied via the upregulation of transporters among which heterodimers SLC7A5/SLC3A2 (LAT1) or SLC7A8/SLC3A2 (LAT2) mediate transport of large neutral amino acids, including Phe.^13^ Without SLC7A5^90^, naïve T cells are unable to proliferate, differentiate or function properly, displaying also profound defects in metabolic reprogramming.^13^ Here, by analysing different cohorts of allergic patients, we observed a reduction of LAT1 and LAT2 in circulating Th2 cells. It might lead to a decrease in intracellular concentration of Phe in memory Th2a subpopulations, which we observed in the most severe cases, paired with an increase of circulating type 2 inflammatory cells (Th2a, CRTH2^+^Tregs and ILC2) and type 2 cytokines. This decrease in LAT1 expression on Th2a cells might result from the increased concentration of Phe in the plasma, as we noted here in patients from different groups, and we confirmed its causality *in vitro*. The physiological concentration of Phe in sera of healthy subjects has been reported to be between 16 to 166μM^91^ with daily variations of 50%^92^ and in phenylketonuria can be greater than 1.2mM.^93^ However, increased systemic levels of Phe and other amino acids in allergic patients^42^ or in animal models of allergic inflammation and helminth infection have been reported previously.^94,95^ Decrease in LAT1 expression in circulating Th2 cells at the beginning of allergic disease may serve as a mechanism preventing excessive activation of Th2 cells, regulating influx of several amino acids, thus limiting local Th2 and ILC2 cell activation.^94,95^ However, in more severe allergic disease, when it is paired with overexpression of other enzymes metabolising Phe, it leads to a decrease of intracellular Phe levels in Th2 cells, which cannot then block OXPHOS, cannot activate IL4I1, or inhibit proliferation which results in unleashed expansion of pathogenic Th2a cells.

### Limitations of the study

This study highlights the pleiotropic effects of Phe metabolism on proliferation, metabolic reprogramming, and function of human memory CD4^+^T cell and Th2 cells and provides description of its impairment in severe allergic diseases in humans. However, it does not deliver mechanistic confirmation in animal models. Although circulating memory CD4^+^T cells often reflect tissue inflammation, as reported by us and others^96, 97, 98^, metabolic assessment of nasal or lung tissue resident memory CD4^+^T cells^45^ would be important to better understand immunometabolic dependencies in local type 2 inflammation. Our metabolomics, proteomics, RNA sequencing, and flow cytometry analyses, while conducted on relatively small cohorts of allergic patients and healthy controls, are mutually validated across each of all studied groups, underscoring the robustness and reliability of our findings. Nonetheless, studies in much larger patient cohorts of different disease severity should follow to further explore the observed phenomena and their utility in prevention and treatment of allergic diseases.

## Supporting information

All supplementary tables

## Acknowledgements and funding

This work was funded by ISCIII (Project numbers, PI19/00044 and PI18/00310) and cofounded by FEDER for the thematic network and co-operative research centres RICORS “Red de Enfermedades Inflamatorias (REI)” (RD21/0002/0008); and the Swiss National Science Foundation (SNSF) grant (nr310030_189334/1) to MS. JRC was supported by an EAACI mid-term research fellowship, CEINDO – BANCO SANTANDER international mobility fellowship and SEMP training fellowship by University of Zurich (UZH). AV acknowledges financial support from “Programa Ayudas Puente 2023-2024” from San Pablo CEU University Foundation and grant number PEJ-2023-AI/SAL-GL-27622 from the Community of Madrid for hiring a research assistant. JCR, NC, JSS and CPT were supported by predoctoral FPIs from Universidad CEU San Pablo. AJGC is supported by grant number PEJ-2023-AI/SAL-GL-27622 from the Community of Madrid, Spain. CG-C was supported by a contract “Atracción de talento investigador” from Community of Madrid, Spain (2016-T2/BMD-1838).

## Contributions

AJK – Conceptualization, Methodology, Investigation, Data analysis, Visualization, Writing-original draft

JRC – Conceptualization, Methodology, Investigation, Data analysis, Visualization, Writing-Review

NS – Investigation, Formal analysis, Visualization, Writing-Review

UR – Investigation, Formal analysis, Visualization, Writing-Review

AJGC – Investigation, Formal analysis, Data analysis, Visualization, Writing-Review

MIDD – Investigation, Formal analysis, Visualization, Writing-Review

NCG – Investigation, Formal analysis, Visualization, Writing-Review IJP – Investigation, Formal analysis, Visualization, Writing-Review

VSS – Conceptualization, Methodology, Investigation, Resources, Writing-Review

PS – Methodology, Writing-Review

EI – Methodology, Writing-Review

CGC – Investigation, Data analysis, Supervision, Writing-Review

JSS – Investigation, Writing-Review

CPT – Investigation, Writing-Review

DO – Investigation, Writing-Review

CMA – Resources, Writing-Review

ME – Resources, Writing-Review

AE – Formal analysis, visualization, Writing-Review

JK – Methodology, Resources, Writing-Review

KB – Methodology, Resources, Writing-Review

AH – Investigation, Writing-Review

GT – Software, Writing-Review

CB – Methodology, Writing-Review

MME – Supervision, Writing-Review and editing, Funding acquisition, Writing-Review

BRL – Resources, Writing-Review

CA – Methodology, Resources, Writing-Review

RJA – Methodology, Resources, Writing-Review

DB – Conceptualization, Supervision, Writing-Review and editing, Project administration, Funding acquisition.

AV – Conceptualization, Methodology, Project administration, Supervision, Writing-Review and Editing

MS – Conceptualization, Methodology, Supervision, Project administration, Writing-original draft, Review and Editing, Funding acquisition

## Declaration of Interests

MS has received research grants from the Swiss National Science Foundation (SNSF nr 310030_189334/1), GSK, Novartis, Stiftung vorm. Bündner Heilstätte Arosa, CK-Care and OM Pharma as well as speaker’s fee from AstraZeneca.

## STAR Methods

## RESOURCE AVAILABILITY

### Lead contact

Further information and requests for resources and reagents should be directed to and will be fulfilled by the lead contact, Milena Sokolowska.

### Materials availability

No new unique reagents were generated in this study.

### Data and code availability

- Metabolomics data have been deposited in National Metabolomics Data Repository (NMDR)/Metabolomics workbench^102^ repository with the accession number ST003135 and are publicly available as of the date of publication. Accession number is also listed in the key resources table.
- All raw data including de-identified human/patient Seahorse, flow cytometry, western blot, SCENITH, qRT-PCR and Olink proteomics data have been deposited as Supplementary Excel tables. They are publicly available as of the date of publication. Accession numbers are listed in the key resources table, when applicable.
- No original code was generated in the study.
- Any additional information required to reanalyze the data reported in this paper is available from the lead contact upon request.

## EXPERIMENTAL MODELS AND STUDY PARTICIPANT DETAILS

### Study participants recruitment Cohort A

The study was approved by the ethical committee of the Hospital Universitario Reina Sofia, Cordoba, Spain (registration number: 5029). Written informed consent was signed by all participants and the study followed all the directives from the Declaration of Helsinki. Study participants were recruited out of the olive pollen season. The Inclusion criteria were: individuals above the age of 18 years, continuously living for a minimum of 10 years in the Cordoba region (one of the areas with a highest seasonal exposure to olive pollen (10.000 pollen grains/m^3^)) and, in case of patients, clinically diagnosed with allergic asthma and/or allergic rhinitis for at least 2 consecutive years along with olive pollen specific IgE (sIgE) ≥ 0.35 KU/L. Lack of signed consent, incomplete understanding of the nature of study, pregnancy and oncological and other inflammatory diseases constituted as the exclusion criteria. Upon recruitment, the participants were divided into 3 groups, namely non allergic controls, mild, and severe allergic patients depending on their sensitization profile and accompanying symptoms^60,61^ as listed:

- Non allergic controls (n=10) – no sensitization to allergens and absence of symptoms
- Mild allergy (n=9) – olive pollen sensitization with IgE against Ole e 7 < 15 kU/L and mild respiratory symptoms
- Severe allergy (n=11) – olive pollen sensitization with IgE against Ole e 7 > 15 kU/L with severe respiratory symptoms during pollen season.

De-identified patients’ demographics and clinical characteristics are available in Tables S9 and S10. Memory CD4^+^Teff and Treg cells from this cohort were analyzed by metabolomics, lipidomics and flow cytometry, whereas serum samples were analyzed by Proximity Extension Assay (PEA) technology (Olink, Uppsala, Sweden), as described below.

### Cohort B

This previously published cohort by Seumois *et. al.,* 2016^68^ comprised of allergic asthma patients (n=37), allergic rhinitis patients (n=25) and healthy controls (n=15). Patients were classified depending on Global Initiative for Asthma (GINA) and skin test reactivity to a total of 32 seasonal and perennial allergens. The study was approved by the Institutional Review Board of La Jolla Institute for Allergy and Immunology, La Jolla, California, USA. Written informed consent was provided by all participants. Participants’ Th2 cells, characterised as CD3^+^CD4^+^CD45RA^−^CD25^−^CCR4^+^ cells were sorted by FACS from peripheral blood mononuclear cells (PBMCs) and their RNA was isolated, amplified and sequenced by HiSeq2500 (Illumina, San Diego, CA, USA). Additional information is available in Seumois *et. al.,* 2016.^68^ RNA-seq data of Th2 cells from this cohort were accessed using GSE75011 and re-analyzed by us, as described below.

### Cohort C

This cohort, previously published by our group, comprised of 16 patients suffering from allergic asthma (mild n=9, severe n=7, classified by GINA) and 8 non-allergic healthy controls with sensitization being defined by Skin Prick Test for aeroallergens. The study was approved by the Committees of Research and Ethics from the Hospital Universitario Puerta de Hierro Majadahonda. Written informed consent was signed by all participants. RNA was isolated from participants’ CD3^+^T cells and subjected to transcriptomics analysis using GeneChip Human Gene 2.1 ST strips (Affymetrix, Thermo Fisher Scientific, Waltham, MA, USA) and manufacturer instructions. Further details can be found in Pablo-Torres *et. al.,* 2023^46^ (GEO: GSE224253). Microarray data from CD3^+^T cells from this cohort were re-analyzed by us, as described below.

### Cohort D

To analyze the systemic level of L-phenylalanine, an additional cohort of olive-pollen allergic patients with the same characteristics as Cohort A was recruited. 8 non-allergic subjects and 67 allergic patients (30 classified as mild and 37 as severe, as in Cohort A) were recruited to analyze L-phenylalanine in serum. The study was approved by the ethical committee of Hospital Clínico San Carlos (date of approval: 19th December 2019; code: 19/552-E_BC). Written informed consent was signed by all participants and the study followed all the directives from the Declaration of Helsinki. Clinical characteristics of this cohort are presented in Table S20. Serum samples from this cohort were analysed by targeted metabolomics, as described below.

### Cohort E

This cohort, previously published by our group^69^, consisted of patients allergic to grass and/or birch pollen (n=13) that underwent allergen immunotherapy, and non-sensitized, non-allergic healthy controls (n=12) which were followed at the same time points. The study was approved by the Ethics Committee of the Canton of Bern (Switzerland). Written informed consent was signed by all participants. The baseline samples, prior to allergen immunotherapy in patients and controls (same time point), were analyzed in the current study. Their allergen sensitization status was concluded using Skin Prick Test and detection of allergen specific IgE. Clinical characteristics of this cohort are presented in Table S21. Further information pertaining to the cohort has been described in Eljaszewicz et. al., 2021.^69^

### Buffy coats and human primary CD4^+^T cell subsets

Buffy coats were obtained from healthy human donors from Chur Blutspendezentrum (Chur, Switzerland) and stored at room temperature prior to processing. Age, gender, and ethnicity of these donors is unknown as these buffy coats were ordered and this information was not requested. Next, PBMCs were isolated using the protocols provided below. Depending on requirements, specific number of PBMCs were further processed while the excess were frozen in freezing medium (90% Heat inactivated Fetal Calf Serum (FCS, Merck, Darmstadt, Germany) + 10% DMSO (Sigma-Aldrich, St. Louis, MO, USA) and stored in liquid nitrogen until future use. Total CD4^+^T, memory CD4^+^T and naïve CD4^+^T cells were always isolated from PBMCs obtained from buffy coats of healthy donors. Total CD4^+^T cells were used in Seahorse Glycolytic Rate and Mito-stress assays and assessment of SLC7A5/LAT1 expression in patient samples. Similarly, human memory CD4^+^T cells were utilized in metabolomics and lipidomics, Seahorse Glycolytic Rate and Mito-stress assays, proliferation and IL4I1 knockdown studies, and flow cytometric analyses. Naïve CD4^+^T cells were isolated from PBMCs and subjected to the Th2 differentiation protocol provided below. Following differentiation, they were used in the qRT-PCR analyses, proliferation, Seahorse ATP-Rate assays, phenotypic characterization, and *SLC7A5*/LAT1 studies.

### *In vitro* differentiation of human Th2 cells

Fresh PBMCs were isolated from buffy coats of healthy donors and resuspended in PBS containing 0.5% BSA (Sigma-Aldrich, St. Louis, MO, USA) and 2mM EDTA (Sigma-Aldrich, St. Louis, MO, USA), as stated previously. Complete RPMI (cRPMI) consists of 10% heat inactivated FCS (Merck, Darmstadt, Germany), 1X Penicillin/Streptomycin (Sigma-Aldrich, St. Louis, MO, USA), 1X Vitamins (Sigma-Aldrich, St. Louis, MO, USA), 1X Sodium pyruvate solution (Sigma-Aldrich, St. Louis, MO, USA), and 1% MEM Non-essential Amino Acid (NEA) (Sigma-Aldrich, St. Louis, MO, USA) solution prepared in RPMI1640 (Lonza, Basel, Switzerland). Naïve CD4^+^T cells were isolated from PBMCs with the autoMACS Pro Separator (Miltenyi Biotec, Bergisch Gladbach, Germany) using the Naïve human CD4^+^T cell isolation kit (Miltenyi Biotec, Bergisch Gladbach, Germany) and resuspended in cRPMI containing recombinant human IL-2 (rhIL-2, 10ng/mL, PeproTech, Thermo Fisher Scientific, Waltham, MA, USA), recombinant human IL-4 (rhIL-4, 12.5ng/mL, PeproTech, Thermo Fisher Scientific, Waltham, MA, USA), anti-IFNγ antibody (5μg/mL, Thermo Fisher Scientific), anti-IL-10 antibody (5μg/mL, Thermo Fisher Scientific) and ImmunoCult Human CD3/CD28 T Cell Activator (25μL/mL, STEMCELL Technologies, Vancouver, Canada) and incubated at 37°C and 5% CO_2_. Cells were maintained at a frequency of 1 million/mL and stimulated every 7 days for a period of 21 days with the activation mix and mentioned antibodies at specified concentrations. cRPMI containing rhIL-2 (10ng/mL, PeproTech, Thermo Fisher Scientific, Waltham, MA, USA), and rhIL-4 (12.5ng/mL, PeproTech, Thermo Fisher Scientific, Waltham, MA, USA) was added between stimulations as per requirement. On day 21, cells were tested for expression of classical Th2 markers, namely, GATA3, CCR4 and CRTH2 by conventional Flow Cytometry to determine their status. Panel provided as in Table S7 and FACSAria III (BD Biosciences, Franklin Lakes, NJ, USA) were used for staining and detection, respectively. Upon confirmation, cells were used for further analyses. This differentiation protocol was adapted from Cousins, Lee, Staynov, 2002^50^. The complete differentiation program is shown in Figure S3.

## METHOD DETAILS

### Isolation of peripheral blood mononuclear cells (PBMCs)

Whole blood was drawn from the study participants into heparin and serum in silica tubes (Vacutainer^®^, BD Biosciences, Franklin Lakes, NJ, USA). Next, PBMCs were isolated using standard Ficoll Paque (Bio & Sell GmbH, Feucht, Germany) by density gradient centrifugation according to the manufacturer instructions. The sera were obtained by centrifugation at 2000g for 10min and collecting the supernatant. Sera and isolated PMBCs were immediately frozen and stored at −80°C until further analysis.

To isolate PBMCs from buffy coats, obtained buffy coats were transferred to sterile flasks and diluted 1:1 in PBS. 30mL of diluted buffy coat was subjected to density gradient centrifugation using Ficoll Paque (Bio & Sell GmbH, Feucht, Germany) and washing steps with cold PBS containing 2mM EDTA. Subsequently, 1X BD Pharm Lyse lysing solution (BD Biosciences, Franklin Lakes, NJ, USA) was used for red blood cell lysis according to manufacturer instructions. PBMCs were washed and resuspended in PBS containing 2mM EDTA for determining counts using trypan blue (Thermo Fisher Scientific, Waltham, MA, USA) and LUNA-II Automated Cell Counter (Logos Biosystems, Gyeonggi-do, South Korea) prior to subsequent processing.

### Fluorescence-activated cell sorting (FACS)

Frozen PBMCs were thawed, washed with warm RPMI1640 medium (Lonza, Basel, Switzerland) and incubated on a gentle roller for 2h at 37°C and 5% CO_2_. Following incubation, cells were washed and stained with eBioscience fixable viability dye eFluor 780 (Thermo Fisher Scientific, Waltham, MA, USA) (1:1000) in the dark at 4°C for 20min. After washing, cells were stained with the antibody panel for T and ILC cells for 30min at room temperature (RT) in the dark. The complete panel of antibodies and isotype controls is presented in Table S1. Cells were then sorted using BD FACSAria III (BD Biosciences, Franklin Lakes, NJ, USA) as follows: Memory CD4^+^Teff cells: Lineage^-^, CD3^+^, CD4^+^, CD8^-^, CD45RA^-^, CD127^+^, CD25^-^; and Memory Treg cells: Lineage^-^, CD3^+^, CD4^+^, CD8^-^, CD45RA^-^, CD127^-^, CD25^+^. Gating strategy used for sorting and subsequent analyses is shown in Figure S1. For metabolomics and lipidomics, cells were sorted to serum free RPMI medium at 4°C. Cells were centrifuged to discard media following which they were immediately fixed with HPLC grade Methanol (MeOH) (Thermo Fisher Scientific, Waltham, MA, USA), following centrifugation at 370g for 5min, and frozen at −80°C until metabolomics analysis was performed. Samples collected from healthy controls (n=6) and allergic patients (n=11) underwent T cell sorting and metabolomics analysis. Samples from additionally recruited participants were analyzed in the same settings, equipment and protocol as described above.

### Metabolomics

#### Sample preparation

As previously published^103,104^, frozen cells in MeOH were thawed at 4°C. Then, a first extraction targeted at lipid metabolites was performed by adding Methyl tert-butyl ether (MTBE) (Sigma-Aldrich, St. Louis, MO, USA) to a final concentration of MTBE:MeOH (1:4). Samples were then sonicated for a total of 15min at 15W (3 rounds of 5min sonication), thoroughly vortexed, and centrifuged for 10min at 16000g to remove any debris. The supernatant was collected. Then a second extraction took place to obtain the most polar metabolites in the sample. To that extent, 100μL of a mixture of H_2_O:MeOH (1:4) was added and the sample was sonicated again for a total of 15min at 15W (3 rounds of 5min sonication), thoroughly vortexed and centrifuged for 10min at 16000g to remove any debris. The supernatant was collected. Polar samples were kept at 4°C until analysis. Extraction solvents were used as a blank and followed the same procedure as the samples. Both, memory CD4^+^T effector and regulatory T cells were analysed as one batch suggesting real biological differences.

Quality control (QC) samples were prepared by pooling equal volumes of samples from the same cell type separately. Then, the QC was aliquoted into different tubes to preserve them from freezing-thawing cycles. Every time a new QC was needed, a fresh QC aliquot was extracted. The QCs clean-up steps followed the same procedure applied to the experimental samples. QCs were analyzed throughout the run to provide a measurement of system stability, performance, and reproducibility of the system.

Additionally, to select features produced only by cells, a calibration curve was made of Teff or Treg cells, respectively. The curve consisted of 4 points that were prepared by seriated dilutions. The number of cells for each point of the Teff cell curve were: 100000, 250000, 500000, and 1500000 cells. For the Treg cells, they were: 10000, 25000, 50000 and 100000 cells. Each cell curve was measured at the beginning, in the middle, and at the end of the analysis of each experimental sample set.

Samples were divided into 2 groups depending on the type of subpopulation they belong to either Teff or Treg. Samples were randomized in 2 consecutive injection sets, analyzing first Treg followed by Teff samples. All samples were measured in duplicate.

#### Analysis by RP-UHPLC-ESI-QTOF-MS analysis

Cell samples were measured using an Agilent HPLC system (1200 series) coupled with quadrupole-time of flight analyzer system (Q-ToF MS 6545) (Agilent Technologies, Waldbronn, Germany).

##### Lipidomics

Data from cell samples were acquired using an Agilent 1290 Infinity II UHPLC system coupled to an Agilent 6550 quadrupole time-of-flight (QTOF) mass spectrometer. The Agilent 1290 Infinity II Multisampler system, equipped with a multi-wash option, was used to uptake 1 and 2µL of extracted samples in positive and negative ionization modes, respectively. The multisampler temperature was maintained at 15°C to preserve lipids in a stable environment and avoid precipitation. An Agilent InfinityLab Poroshell 120 EC C18 (3.0 × 100mm, 2.7µm) (Agilent Technologies, Waldbronn, Germany) column and a compatible guard column (Agilent InfinityLab Poroshell 120 EC C18, 3.0 × 5mm, 2.7µm) were used and maintained at 50°C. The chromatography gradient started at 70% of B at 0 – 1min, 86% at 3.5 – 10min, 100% B at 11–17min. The starting conditions were recovered by minute 17, followed by a 2min re-equilibration time; the total running time was 19min. The mobile phases used for both positive and negative ionization modes consisted of (A) 10mM ammonium acetate, 0.2mM ammonium fluoride in 9:1 water/methanol and (B) 10mM ammonium acetate, 0.2mM ammonium fluoride in 2:3:5 acetonitrile/methanol/isopropanol. The flow rate was held constant, set at 0.6mL/min. The multi-wash strategy consisted of a mixture of methanol:isopropanol (50:50, v/v) with the wash time set at 15s, and aqueous phase:organic phase (30:70, v/v) mixture to assist in the starting conditions.

The Agilent 6550 QTOF mass spectrometer equipped with a dual AJS ESI ion source was set with the following parameters: 150V fragmentor, 65V skimmer, 3500V capillary voltage, 750V octopole radio frequency voltage, 10L/min nebulizer gas flow, 200°C gas temperature, 50psi nebulizer gas pressure, 12L/min sheath gas flow, and 300°C sheath gas temperature. Data were collected in centroid in positive and negative ESI modes in separate runs, operated in full scan mode from 50 to 1700*m/z* with a scan rate of 3 spectra/s. A solution consisting of two reference mass compounds was used throughout the whole analysis: purine (C_5_H_4_N_4_) at *m/z* 121.0509 for the positive and *m/z* 119.0363 for the negative ionization modes; and HP-0921 (C_18_H_18_O_6_N_3_P_3_F_24_) at *m/z* 922.0098 for the positive and *m/z* 980.0163 (HP-0921+acetate) for the negative ionization modes. These masses were continuously infused into the system through an Agilent 1260 Iso Pump at a 1mL/min (split ratio 1:100) to provide a constant mass correction. Ten Iterative-MS/MS runs were performed for both ion modes at the end of the analytical run. They were operated with an MS and MS/MS scan rates of 3spectra/s, 40– 1700*m/z* mass window, a narrow (∼ 1.3amu) MS/MS isolation width, 3 precursors per cycle, 5000 counts, and 0.001% of MS/MS threshold. Five iterative-MS/MS runs were set with a collision energy of 20eV, and the subsequent five runs were performed at 40eV. References masses and contaminants detected in blank samples were excluded from the analysis to avoid inclusion in the iterative-MS/MS.

Additionally, confirmation of some compounds was performed by LC-MS/MS experiments using 20V for fragmentation.

##### Polar extraction metabolomics

The HPLC system was equipped with a degasser, two binary pumps, and a thermostated autosampler. Briefly, 2μL of the sample were injected into an Agilent Zorbax Extend C18 (2.1 × 50mm, 1.8μm; Agilent, Waldbronn, Germany), with a guard column Zorbax Extend C18 (3 × 5mm, 1.8μm; Agilent, Waldbronn, Germany), both maintained at 60°C. The flow rate was set at 0.6mL/min. The elution gradient involved a mobile phase consisting of: (A) water containing 0.1% of formic acid and (B) acetonitrile containing 0.1% of formic acid. The initial conditions were set at 5% phase B for 1min, which increased linearly to 80% phase B in 7min. Then in 4.5min it increased until 100% of phase. Then the equipment returned to the initial condition in 0.5min, which were held for 3min for column reconditioning. Samples were analyzed in both ESI+ and ESI-modes in separate injections. The capillary voltage was set at 3000 for both polarities. The drying gas flow rate was 12L/min at 250°C and gas nebulizer at 52psi; fragmentor voltage was set at 175V in ESI+ and 250 in ESI-; skimmer and octupole radio frequency voltages were set to 65 and 750V, respectively. MS spectra were collected in the centroid mode at a scan rate of 3spectra/s. The MS detection window was performed in a full scan from 100 to 1200*m/z* for both modes. Automatic MS recalibration during batch analysis was carried out by introducing a reference standard into the source via a reference sprayer valve. Reference masses for ESI+ were purine (*m/z* = 121.0508) and HP-0921 (*m/z* = 922.0097), whereas for ESI− TFA NH_4_ (*m/z* = 112.9855) and HP-0921 (*m/z* = 966.0007).

#### Data processing

Memory CD4^+^Teff and Treg raw data were re-processed together so they could be compared, as each feature has an average mass and an average retention time (unique ID in both cell types). The data collected after the LC-MS analyses in both positive and negative ion modes were reprocessed with the Agilent MassHunter Profinder B.10.0.2 software. The datasets were extracted using the Batch Recursive Feature Extraction (RFE) workflow integrated into the software. This workflow comprises two steps: the Batch Molecular Feature Extraction (MFE) and the Batch Find by Ion Feature extraction (FbI). The MFE algorithm consists in removing unwanted information, including the background noise, and then creating a list of possible components that represent the full range of TOF mass spectral data features, which are the sum of coeluting ions that are related by charge-state envelope, isotope pattern, and/or the presence of different adducts and dimers. Additionally, in the case of lipidomics, the MFE is intended to detect coeluting adducts of the same feature, selecting the following adducts: [M+H]^+^, [M+Na]^+^, [M+K]^+^, [M+NH4]^+^ and [M+C2H6N2+H]^+^ in LC-MS positive ionization; [M-H]^−^, [M+CH3COOH-H]^−^, and [M+Cl]^−^ in LC-MS negative ion mode. Whereas, in polar analysis, the following adducts: [M+H]^+^, [M+Na]^+^, [M+K]^+^, and [M+NH4]^+^ in LC-MS positive ionization; [M-H]^−^, [M+HCOOH-H]^−^, and [M+Cl]^−^ in LC-MS negative were selected. The neutral loss (NL) of water is also considered for both ion modes in the two methods (lipidomics and polar analysis). The algorithm then aligns the molecular features across the study samples using the mass and retention time (RT) to build a single spectrum for each compound group. The next step involves FbI, using the median values derived from the MFE process to perform a targeted extraction to improve the reliability of finding and reporting features from complex datasets used for differential analysis.

After data pre-processing, the quality assurance (QA) of the datasets were reprocessed independently according to their blank samples, experimental samples, QC samples, and their specific cell calibration curve. This consisted of raw data filtration by keeping all features that were present after blank subtraction and were detected in >75% of QCs and >75% in at least one sample group (A (controls), B (mild allergy), C (severe allergy)) and had Relative Standard Deviation (RSD) <30% in the QCs. The rest of the signals were excluded from the analyses. Finally, features were kept if they had a significant correlation (ρ) with the number of cells in the curve (|ρ| > 0.6 and p value < 0.05), leading to 109 and 190 chemical entities were obtained that passed LC-MS quality control for lipidomics and polar analysis, respectively. It is important to note that some of the metabolites that are present only in one of the cell subsets might also be present in the other, and the absence only indicates that it does not comply with the QA.

Missing values were replaced using the k-nearest neighbors (kNN) algorithm^105^ using an in-house script developed in Matlab® (v.R2018b, MathWorks®). Data were normalized using the number of cells obtained from the cell sorter. The quality of the analyses was tested using principal component analysis (PCA) models^106^.

#### Workflow for lipid annotation

The annotation process was carried out in three steps: the first one was an initial tentative identification of lipid features, based on the MS1 data, using our online tool CEU Mass Mediator (CEUMM) (http://ceumass.eps.uspceu.es/mediator/)^104,107^. This tool for mass-based compound annotation comprises the information available in different databases (KEGG, HMDB, LIPID MAPS, Metlin, MINE, and an in-house library). This stage started with the tentative assignment based on (i) accurate mass (maximum mass error tolerance 20 ppm); (ii) RT; (iii) isotopic pattern distribution; (iv) the possibility of cation and anion formation; and (v) adduct formation pattern.

Secondly, to increase the level of annotation confidence, the raw LC-MS/MS data from lipidomics was imported to a Lipid Annotator software (Agilent Technologies, Waldbronn, Germany) to build a fragmentation-based (MS/MS) library comprising the *m/z* of all the precursors identified as lipids by the software, together with their corresponding RT. The Lipid Annotator method^108^ was set as follows: ion species [M+H]^+^, [M+Na]^+^, and [M+NH4]^+^ for positive; and [M-H]^−^, and [M+CH3COOH-H]^−^ for negative ionization mode. Then, for both ion modes, the Q-Score was set at ≥ 50; all the lipid classes were selected, the mass deviation was established as ≤20ppm, fragment score threshold was fixed as ≥30, and the total score was set at ≥60. Additionally, MS-DIAL software^109^ was used for lipidomics and polar analysis iteratives. For this software, analytical parameters were set as described: ion species [M+H]^+^, [M-H2O+H]^+^, [M+Na]^+^, [M+K]^+^, [M+NH4]^+^ and [M+C2H6N2+H]^+^ in positive ionization mode; and [M-H]^−^, [M-H2O-H]^−^, [M+Cl]^−^, [M+HCOOH-H]^−^ and [M+CH3COOH-H]^−^ in negative mode for lipidomics. Adduct formation with formic acid was observed experimentally in this study even though the mobile phases used for LC are lacking this compound. Formic acid presence was considered due to trace amount contamination levels of formate in acetate salts of LC-MS grade, or methanol oxidation during ESI (https://lipidomicstandards.org/isomeric-overlap/). In the case of polar analysis, ion species [M+H]^+^, [M-H2O+H]^+^, [M+Na]^+^, [M+K]^+^ and [M+NH4]^+^ in positive mode; and [M-H]^−^, [M-H2O-H]^−^, [M+Cl]^−^, and [M+HCOOH-H]^−^ in negative were selected. Regardless of the analytical method or ionization mode, the search was fixed to be performed across a mass range from 50 to 1500Da (for both MS1 and MS/MS levels), the mass deviation accepted was ≤0.01Da and ≤0.025 for MS1 and MS/MS levels respectively, and the identification score cut off was also set at ≥60. Metabolites identification was performed against inside MS-DIAL lipidomic database^109^ and MassBank of North America (MoNA) database for polar compounds (https://mona.fiehnlab.ucdavis.edu/downloads). Finally, for targeted LC-MS/MS experiments, manual spectral interpretation^110^ was carried out using the software Agilent MassHunter (version 10.0), matching the retention time and MS/MS fragmentation to the available spectral data included in MetFrag and Lipid Maps.

Metabolomics data have been deposited in National Metabolomics Data Repository (NMDR)/Metabolomics workbench^102^ repository with the accession number ST003135 and they are also available in Table S2.

### Targeted analysis of Phenylalanine

Phenylalanine concentration was analysed in Cohort D using an adaptation of the previously published method^101^. In brief, serum samples were prepared in a randomized order and measured in batches using dynamic molecular reaction monitoring on a liquid chromatography system (1260 Infinity II, Agilent Technologies, Waldbronn, Germany) coupled to a triple quadrupole mass spectrometer with electrospray ionization Agilent Jet Stream source, 6470 Agilent Technologies.

To analyze Phenylalanine, a gradient elution on a Kinetex HILIC (150mm × 2.1mm × 100Å) column maintained at 25°C was used. The mobile phases were (A) water, and (B) Acetonitrile, both with 5.5mM ammonium acetate and 0.1% acetic acid. The flow rate was 0.5mL/min, and the gradient started with 5% of A for 2min, increased up to 50% until 12min, and went back to initial conditions until 22min. The MS conditions were: 5500V of capillary voltage in positive ESI mode, a nebulizer gas flow rate of 11.0L/min, a source temperature of 250°C; and a source pressure of 60psi.

For sample preparation, 50µL of serum were mixed with 150µL of cold (−20°C) methanol: ethanol mix (MeOH:EtOH) (1:1). Then, serum samples were vortex-mixed for 20s, placed on ice for 20min and centrifuged at 16000g for 20min at 4°C. Then, 70µL of the supernatant was transferred into an LC vial and mixed with 50µL of ISTD mix] ^101^ and 440µL of the initial conditions of the mobile phases (95% B + 5% A). QC and blank samples, prepared as described above, were also measured through the run.

### Flow cytometry data analyses

Flow cytometry data were analyzed with classical gating and two-dimensional reduction using t-distributed stochastic neighbor embedding (tSNE) algorithms using FlowJo X v.10.7.1. (BD Biosciences, Franklin Lakes, NJ, USA). First, the data were cleaned by using the FlowClean algorithm^111^. Next, classical gating was applied for T cell populations and ILCs (for representative gating strategy, please see Figure S13). For two-dimensional reduction, memory CD4^+^Teff cells (CD45RA^-^CD3^+^CD4^+^CD127^++^CD25^-^) were gated and downsampled using DownSample v3.3.1. to ensure that analysis are performed with the same number of events, namely, the highest event number in the available gate. Next, all files were concatenated into a new data file (separate for allergic individuals and controls). Finally, the FlowJo tSNE algorithm^112^ was applied (Iterations: 1000 Perplexity: 30 Eta: 9100 KNN: Exact, Gradient algorithm: Barnes-Hut) and analyzed with FlowSOM v4.0.^113^ and Cluster Profiler 1.7.4. based on CD161, CRTH2, cKit, CD127, CD25 and PD1 expression. The populations obtained with FlowSOM were back-gated in the original files to obtain the total frequency of the population of interest in each sample.

### Serum Protein Extension Assay (PEA) proteomics and targeted immunoassays

Protein expression in the sera of patients from Cohort A was measured using the human Protein Extension Assay technology (Olink, Uppsala, Sweden). Olink Target 96 Immuno-Oncology, Cardiovascular II and Cardiometabolic panels were used according to manufacturer’s instructions. Olink data in normalized protein expression (NPX) format were imported, processed, and compared between groups with OlinkR package (https://github.com/ge11232002/OlinkR). A protein was considered expressed if it was above the detection limit in more than half of the samples in either of the two compared groups.

Hepatic growth factor (HGF) was measured following manufacturers’ instruction in serum samples diluted 1:2 in PBS. The plate was analyzed on LUMINEX 200 (Luminex Corporation, Austin, TX, USA) and Bio-Plex Manager 6.1 (Bio-Rad Laboratories, Hercules, CA, USA).

### Seahorse Flux Analyses

For Seahorse flux analysis, fresh PMBCs were isolated from buffy coats as described previously. Next, total CD3^+^CD4^+^T cells or memory CD3^+^CD4^+^T cells were isolated using the human CD4^+^T Cell Isolation Kit or Memory CD4^+^T Cell Isolation Kit (Miltenyi Biotec, Bergisch Gladbach, Germany), for respective cell types, according to manufacturer instructions on the autoMACS Pro Separator (Miltenyi Biotec, Bergisch Gladbach, Germany). Isolated cells were then transferred to RPMI1640 medium (Lonza, Basel, Switzerland) with 10% heat inactivated FCS (Merck, Darmstadt, Germany), 1% NEA (Merck, Darmstadt, Germany) and 100U/mL rhIL-2 (PeproTech, Thermo Fisher Scientific, Waltham, MA, USA) (R10+IL2). Cells were then incubated for 24, 48 or 72h in this culture medium additionally supplemented with either 0.1mM or 1mM Arg or Phe or Vehicle (Merck, Darmstadt, Germany). After incubation, cells where washed and their energy metabolism was assessed using Glycolytic rate and Mito stress Seahorse flux analysis assays (Agilent Technologies, Santa Clara, CA, USA) according to manufacturer instructions. Acute activation of T cells was performed directly during these assays, before the injection of any inhibitor, using anti-CD2, anti-CD3, and anti-CD28 antibody coated beads (Miltenyi Biotec, Bergisch Gladbach, Germany) at a ratio of 3:1 (beads:cells). A similar protocol was used for *in vitro* differentiated Th2 cells wherein cells were incubated in appropriate culture medium for 24h, washed, and seeded in a Seahorse plate pre-coated with Poly-D-Lysine (50μg/mL) at a concentration of 1×10^5^ cells/well, after which Seahore ATP rate assay was performed as per manufacturer instructions. Following completion of the assay, the supernatant in each well was discarded and the plate was frozen at −80°C for normalization purposes. Data normalization was done either by quantifying the total protein content using the Pierce BCA protein quantification assay (Thermo Fisher Scientific, Waltham, MA, USA) by manufacturer instructions or by cell count since they were seeded on the day of analysis. Preliminary data analysis was performed on the Seahorse Wave Desktop Software (Agilent Technologies, Santa Clara, CA, USA) following which data were exported and further analysis was done using Prism (GraphPad Software Inc., Boston, MA, USA)

### CFSE based Proliferation assay

Total CD3^+^CD4^+^T cells or memory CD3^+^CD4^+^T cells were isolated from buffy coats as described before and cultured in R10+IL2. Cells were stained with 5μM or 10μM CFSE (Thermo Fisher Scientific, Waltham, MA, USA) following manufacturer instructions and seeded at a concentration of 10^6^/mL in R10+IL2 supplemented additionally with 0.1mM, 1 mM of Phe (Merck, Darmstadt, Germany) or Vehicle and stimulated with anti-CD2, anti-CD3 and anti-CD28 antibodies (Sigma Aldrich, St. Louis, MO, USA) and incubated at 37°C and 5% CO_2_. Every 24h, cells were harvested and stained with 1μL of eBioscience viability dye eFluor 780 (Thermo Fisher Scientific, Waltham, MA, USA) in the dark at 4°C for 20min and acquired on a BD Fortessa flow cytometer (BD Biosciences, Franklin Lakes, NJ, USA). Proliferation of *in vitro* differentiated Th2 cells was also assessed using the same protocol and data was acquired on BD FACSAria III (BD Biosciences, Franklin Lakes, NJ, USA).

### Quantitative Reverse Transcription-Polymerase Chain Reaction (qRT-PCR)

Memory CD3^+^CD4^+^T cells were isolated, cultured and treated with Phe or Vehicle as mentioned above. *In vitro* differentiated Th2 cells were incubated in R10+IL2 supplemented with 0.05mM, 0.1mM, 0.5mM and 1mM Phe or Vehicle with or without bead-based T cell stimulation for 24h. The base medium contains approximately 1.14mM of L-arginine and 90.9μM of L-phenylalanine. The final concentration of phenylalanine in these media are 0.1454mM, 0.1909mM, 0.59mM and 1.09mM, respectively. Similarly, the final concentration following supplementation of 0.05mM, 0.1mM, 0.5mM and 1mM Arg are 1.18mM, 1.23mM, 1.63mM and 2.13mM, respectively. Cells were harvested and lysed on ice with RLT buffer (QIAGEN, Hilden, Germany) containing β-mercaptoethanol (10μL/mL of RLT buffer) (Sigma Aldrich, St. Louis, MO, USA), and stored at −80°C until analyses. mRNA isolation was performed using RNeasy Micro or Mini kits (QIAGEN, Hilden, Germany) according to manufacturer’s instructions. Quality and quantity of isolated RNA was assessed by Nanodrop 2000 (Thermo Fisher Scientific, Waltham, USA). The flow-through from the RNeasy spin column was saved and stored at −80°C for protein precipitation. Reverse transcription was performed with RevertAid RT kit (Thermo Fisher Scientific, Waltham, MA, USA) and random hexamers according to the manufacturer’s recommendations. Gene expression (5ng cDNA per well) was assessed by qRT-PCR using 5μM of appropriate primers and SYBR Green/ROX qPCR Master Mix (Thermo Fisher Scientific, Waltham, MA, USA), performed on QuantStudio 7 Flex Real-Time PCR System (Thermo Fisher Scientific, Waltham, MA, USA). Genes expression was normalized to either elongation factor 1α (EEFA1) or 18S rRNA and presented as relative quantification calculated with the ΔΔC_t_ formula using the Vehicle-treated cells as a calibrator. A complete list of the primers used is available in the key resources table.

### IL4I1 knockdown in human memory CD4^+^T cells and proliferation

IL4I1 knockdown was carried out using siRNA methodology and Human T Cell Nucleofector Kit (Lonza, Basel, Switzerland) using manufacturer instructions. Memory CD4^+^T cells from PBMCs of healthy donors were isolated using MACS technology as described previously and allowed to rest overnight in R10+IL2 at 37°C and 5% CO_2_. On the next day, 5 million cells were washed and resuspended in 100μL of prepared nucleofection reagent, and SMARTPOOL ON-TARGETplus human IL4I1 siRNA or Negative Control siRNA (final concentration 300nM) (Horizon Discovery, Cambridge, UK) and added to the supplied cuvettes and subjected to electroporation using the U-14 program on Amaxa Nucleofector I device (Lonza, Basel, Switzerland). Following electroporation, cells were incubated in warm RPMI1640 for 4h at 37°C and 5% CO_2_ and then transferred to R10+IL2 and incubated overnight at 37°C and 5% CO_2_. Post incubation, cells were washed and stained with 5μM CFSE (Thermo Fisher Scientific, Waltham, MA, USA) according to manufacturer’s instructions. Cells were then transferred to media supplemented with 1mM Phe or Vehicle with (1:1 cells:beads) or without anti-CD2, anti-CD3 and anti-CD28 antibody coated bead (Merck, Darmstadt, Germany) based stimulation for 24 or 48h. At both time points, cells were harvested and stained with BD Horizon Fixable Viability Stain 780 (1:1000) (BD Biosciences, Franklin Lakes, NJ, USA) for 15min at RT in the dark following which they were stained with BV510 conjugated anti-CD45 antibody (1:100, clone H130, BioLegend, San Diego, CA, USA) and incubated for 30min at 4°C in the dark. Cells were then washed, and data was acquired on BD FACS Aria III (BD Biosciences, Franklin Lakes, NJ, USA).

### SCENITH: Single Cell ENergetIc metabolism by profilIng Translation inHibition

SCENITH is a flow cytometry-based technique used to assess cellular metabolism, developed by Arguello *et. al.,* 2020^49^. Frozen PBMCs were thawed, resuspended in RPMI1640 containing 10% FBS and 1X NEA and allowed to recover for a minimum of 2h. After incubation, PBMCs were washed and added to R10+IL2 supplemented with 1mM Phe or Vehicle and incubated for 12-14h. Post this incubation, cells were harvested and subjected to the SCENITH protocol^49^. In short, cells were treated with Control, 2-DG, Oligomycin, 2-DG and Oligomycin and Harringtonine inhibitors at recommended dilution for 15min at 37°C. Puromycin was added following incubation to block protein translation and incubated at 37°C for 30min. Following incubation, cells were first washed and stained with Fixable Viability stain 780 (1:1000, BD Biosciences, Franklin Lakes, NJ, USA) for 15min in the dark. After incubation, they were stained with a panel of antibodies specific for Th2 and Treg surface markers (Table S8) for 30min at 4°C. Cells were washed subjected to intranuclear staining using the FOXP3 intranuclear staining kit (Thermo Fisher Scientific, Waltham, MA, USA) to detect levels of puromycin incorporation along with intranuclear flow antibodies. Samples were then washed and read on the BD FACS Aria III and analysed according to formulas mentioned in the original publication^49^.

### RNA-seq and microarray data analysis

RNA-seq data of Th2 cells and microarray data of total CD3^+^T cells were downloaded from the publicly available Gene Expression Omnibus platform under accession numbers GSE75011 (Cohort B) and GSE224253 (Cohort C), respectively. Data were analyzed with the GEO2R interactive web tool available here: www.ncbi.nlm.nih.gov/geo/geo2r/. Enrichment analysis of the most significant process networks in Th2 cells from patients with allergic asthma and allergic rhinitis, when compared to Th2 cells of non-allergic individuals, was performed with Metacore software version 23.4.71500 (Thomson Reuters, Toronto, Canada) using enrichment by GO Processess (all significantly changed genes) and by Metabolic Networks (significantly upregulated and significantly downregulated genes) based on p-value<0.05 (Tables S13-S18). The phenylalanine pathway gene set was curated and adapted from GSEA and MSigDB Database (Broad Institute, MIT and Reagent of the University of California, USA) (Table S19) available under systematic names: M29207, M37685, M46134, M16650, M27835, M27850, M45537. Only genes present in the Th2 RNA-seq dataset analysis were shown in Figures 7D and S14D. The same genes were followed in our previously published CD3^+^T cell microarray dataset and are shown in Figures 14E and 14F.

### Western Blotting

The flow through from the RNeasy spin column during RNA isolation, saved and stored at −80°C, was thawed and processed according to the manufacturer’s instructions to obtain total protein. Briefly, 4 volumes of ice-cold acetone (Sigma-Aldrich, St. Louis, MO, USA) was added to 1 volume of flow through and incubated at −20°C for 30min. After incubation, the mix was centrifuged at maximum speed for 10min at 4°C in a benchtop ultracentrifuge and the supernatant was discarded. The pellet was washed with 100μL of ice-cold ethanol, centrifuged, and allowed to air dry after the supernatant was discarded. Then, the pellet was resuspended in 45μL 1X Laemlli buffer (270μL of 4X Laemmli buffer, 30μL of β-mercatopethanol, 900μL of water) and heated at 95°C for 5min. After heating, the samples were cooled and loaded onto a SurePAGE Bis-Tris 4-12% gel (GenScript, Piscataway, NJ, USA) with Protein Precision Plus dual color ladder (Bio-Rad laboratories, Hercules, CA, USA) and subjected to electrophoresis at 100V. Then, by means of the eBlot L1 Protein Transfer System (Genscript, Piscataway, NJ, USA), proteins were transferred from the gel onto a prepared WesternBright NC nitrocellulose membrane (Advansta Inc., San Jose, CA, USA) as per supplier instructions. The membrane was washed and blocked for 2h at room temperature with buffer containing 5% milk, and 0.05% Tween-20 prepared in PBS. Following blocking, we probed the membrane with either Rabbit anti-LAT1 (1:1000, Cell Signaling Technologies, Danvers, MA, USA) or Rabbit anti-IL4I1 antibody (1:1000, Thermo Fisher Scientific, Waltham, MA, USA) prepared in blocking buffer and incubated it overnight at 4°C. After incubation, the membrane was washed with PBS containing 0.05% Tween 20 (PBS-T) and probed with HRP conjugated goat anti-rabbit IgG (H+L) antibody (1:10000, Jackson Laboratories, Bar Harbor, ME, USA) for 1h at RT. The membrane was washed and developed with WesternBright Quantum reagent (Advansta Corporation, Menlo Park, CA, USA) per manufacturer instructions and read on Fusion FX 7 imaging system (Vilber Lourmat, Eberhardzell, Germany). The membrane was stripped with Restore PLUS Western Blot Stripping buffer (Thermo Fisher Scientific, Waltham, MA, USA), washed, blocked and probed with anti-human beta actin antibody (1:1000, Thermo Fisher Scientific, Waltham, MA, USA) and processed with the same steps. Relative protein expression of LAT1 and IL4I1 were calculated using beta-actin as a calibrator based on their intensity levels determined using Fiji 2.14.0/1.54f (NIH, Bethesda, MD, USA).

### Phenylalanine quantification in *in vitro* experiments

*In vitro* differentiated Th2 cells were transferred to R10+IL2 supplemented with additional 1mM Phe or Vehicle. Importantly, 10μM of LAT1 inhibitor KYT0353 (Tocris Bioscience, Avonmouth, UK) was also added for selective inhibition and incubated for 6h. Upon completion of incubation, cells were harvested and immediately lysed in the Phenylalanine Assay buffer as per Phenylalanine Assay kit (Abcam, Cambridge, UK) instructions which was used to quantify Phe levels in cell lysates. Samples were processed according to manufacturer instructions and OD was measured at 450nm on the Mithras LB 940 plate reader (Berthold Technologies, Bad Wildbad, Germany) after incubation. Phe quantification in these samples and subsequent data analysis and calculations were carried out by following manufacturer instructions.

## QUANTIFICATION AND STATISTICAL ANALYSIS

### Statistical analysis

Multivariate analysis was performed using SIMCA v.16.0 (Sartorius Stedim Data Analytics). The PCA models were used to evaluate data quality and find patterns in the experimental samples. The cross-validated Orthogonal Partial Least Square Discriminant Analysis (OPLS-DA) models were performed to confirm the separation between the metabolism of memory CD4^+^Teff and Tregs. The models were evaluated using R^2^ and Q^2^ parameters which are the classification and prediction capacity, respectively.

Complementary, univariate analysis was performed in MATLAB (v.R2018b, MathWorks®) to obtain the significance of each feature in the study between studied groups. Thus, non-parametric Mann-Whitney U was used to determine statistical significance between pair groups, and the statistical significance was set at p-value<0.05. Additionally, for multiple comparison correction, False Discovery Rate (FDR) was performed by using Benjamini-Hochberg correction for these p-values, and statistical significance was set as well at FDR<0.05 for the adjusted p-value^114^. Bar representations were obtained in GraphPad Prism version 9.0.0.

Venny online tool (v. 2.0) was used to construct the Venn diagrams; and the MetaboAnalyst online tool (v. 5.0) was used to produce heatmaps graphs where data was logarithmic transformed and auto scaled. When hierarchical clustering was applied, Euclidean distance measure and Ward’s clustering method were chosen as the clustering parameters. Moreover, enrichment pathway analysis was performed with IMPaLA pathway over-representation analysis tool. Two analyses, one for Teff cells and Treg cells each, were performed using those identified metabolites with an associated HMDB ID number found in both and in the specific cell subset (n = 48 metabolites for Teff cells and n = 51 for Treg cells). Only routes from KEGG database were considered.

For comparisons between two groups, t-tests either unpaired or paired t-test, or Mann-Whitney U or Wilcoxon tests were used depending on data distribution and variable. Similarly, One-way ANOVA with Fisher’s LSD, or Tukey, or Sidak’s, or Dunnett correction were used for comparison of 3 or more groups depending on distribution and variable type. For all data analysis, sample sizes and appropriate analysis parameters including tests used are indicated in respective figure legends. GraphPad Prism v10.2.0 was used for analysis. Significance was concluded at *p< 0.05, **p< 0.01, ***p< 0.001, ****p<0.0001.

