## Supplementary figures, legends and table legends for "L-Phenylalanine is a metabolic checkpoint of human Th2 cells"

Figure S1

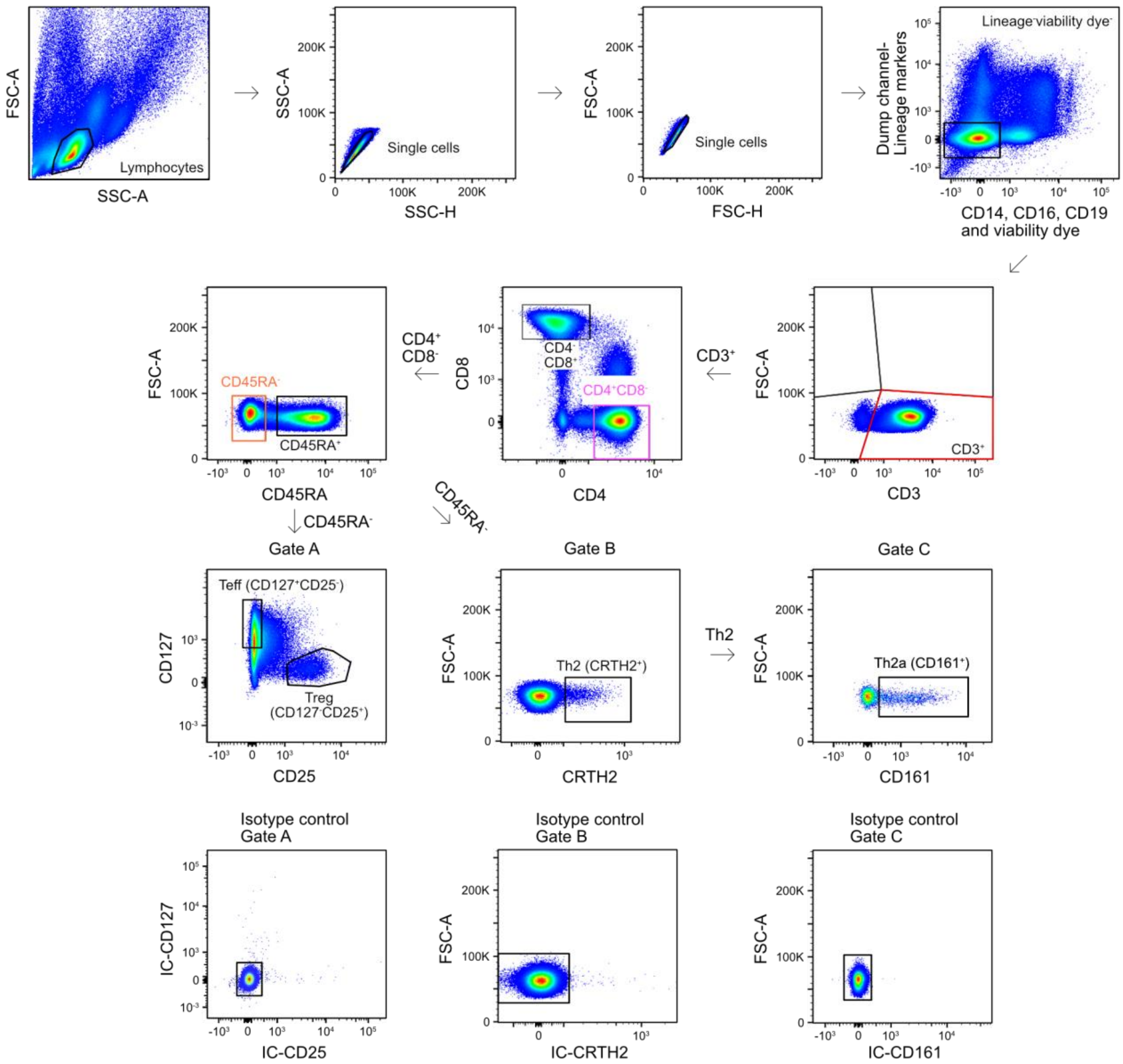


**Figure S1. Gating strategy for T cells used for sorting circulating human memory CD4⁺Teff and Treg cells and classical analysis.** Briefly, T cells were gated by eliminating doublets and negatively selecting dead and other cells positive for lineage specific markers. $CD3^+$, $CD4^+$, $CD8^-$ cells were positively gated. Subsequently, memory Teff ($CD45RA^-$, $CD127^+$, $CD25^-$) and memory Treg ($CD45RA^-$, $CD127^-$, $CD25^+$) cells were sorted using gate A. Th2 and Th2a (“pathogenic”) cells were determined by the presence of CRTH2 (gate B) and double expression of CRTH2 and CD161 (gate C), respectively. Respective isotype controls are shown.

Figure S2

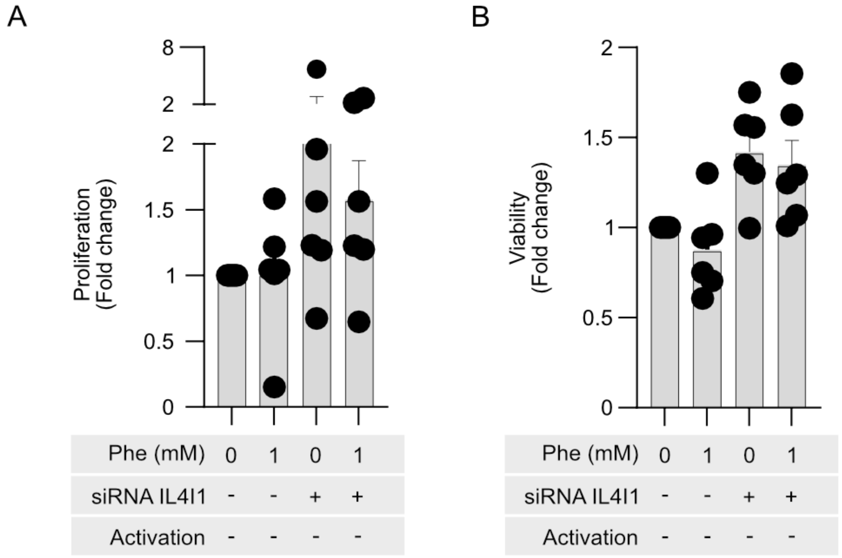


**Figure S2. Proliferation of non-activated memory CD4⁺T cells following IL4I1 knockdown. (A)** Quantification of proliferation of control siRNA (Crtl) and siRNA IL4I1-treated human memory CD4⁺T cells from 2 independent experiments in 5 different donors, incubated in full medium with/without 1mM of additional Phe and without CD2, CD3 and CD28 activation antibodies treatment for 24h prior to flow cytometry. **(B)** Quantification of viability of siRNA Crtl and siRNA IL4I1-treated human memory CD4⁺T cells by flow cytometry in the same experiments as in (A). Bar graphs show a fold change in proliferation as compared to the Vehicle-treated and activated cells. Wilcoxon test was used for analysis. All data are presented as mean±SEM. *p<0.05, **p<0.01, ***p<0.001. Ctrl, control

Figure S3

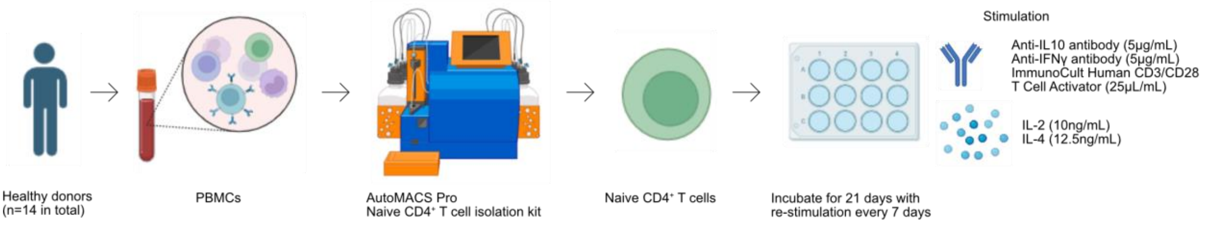


**Figure S3. Methodology and timeline used for *in vitro* differentiation of naïve CD4⁺T cells into Th2 cells.** Briefly, naïve CD4⁺T cells from PBMCs of healthy subjects are isolated and incubated in media containing IL-2 (10ng/mL), IL-4 (12.5ng/mL), anti-IFNγ antibody (5μg/mL), anti-IL-10 antibody (5μg/mL) and ImmunoCult Human CD3/CD28 T Cell Activator (25μL/mL) for 21 days with restimulation every 7 days. Further details are provided in STAR Methods. This differentiation protocol was adapted from Cousins, Lee, Staynov, 2002[50]. Prepared with Biorender.com.

Figure S4

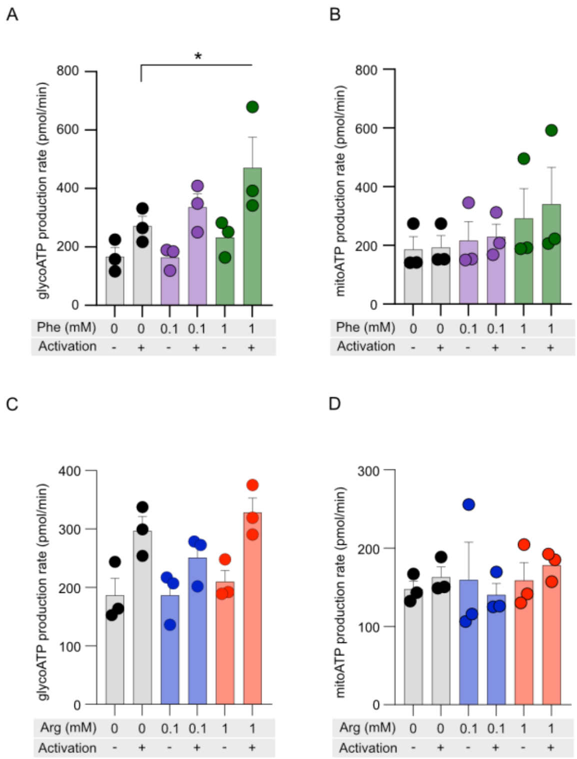


**Figure S4. Quantification of induced glycoATP and mitoATP rates upon L-phenylalanine and L-arginine supplementation in *in vitro* differentiated Th2 cells.** (**A-B**) Quantification of induced glycoATP rate (**A**) and induced mitoATP rates (**B**) of *in vitro* differentiated Th2 cells incubated in full culture medium with/without additional Phe at concentrations of 0.1mM, 1mM, or Vehicle for 24h with/without acute CD2, CD3 and CD28 antibodies activation. (**C-D**) Quantification of induced glycoATP rate (**C**) and induced mitoATP rates (**D**) of *in vitro* differentiated Th2 cells incubated in full culture medium with/without additional Arg at concentrations of 0.1mM, 1mM, or Vehicle for 24h with/without acute CD2, CD3 and CD28 antibodies activation. Bars represent mean±SEM. Three independent experiments with 3 donors were conducted. Each dot represents the mean of more than 4 technical replicates per donor (n=3 different donors). One-way ANOVA with Sidaks´s multiple comparison correction was used to assess statistical significance. *$p<0.05$.

Figure S5

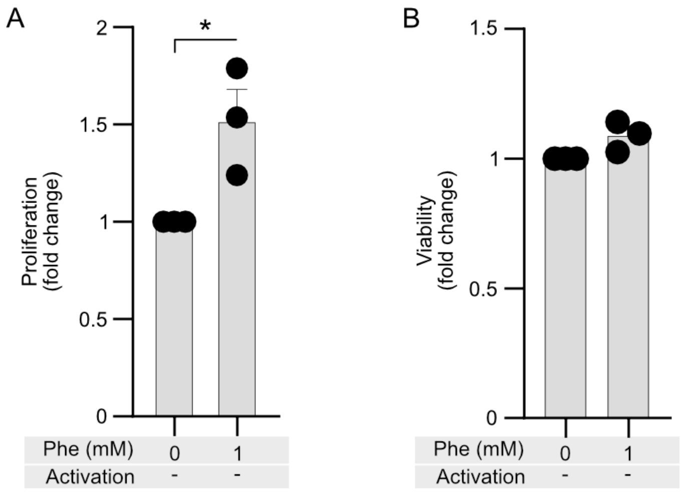


**Figure S5. Effect of L-phenylalanine on the proliferation of non-activated *in vitro* differentiated Th2 cells.** (**A-B**) Proliferation (**A**) and viability (**B**) of *in vitro* differentiated Th2 cells subjected to high doses of Phe. *In vitro* differentiated Th2 cells were incubated in full culture medium with or without additional supplementation of 1mM Phe without activation with CD2, CD3 and CD28 antibodies for 24h. Then, proliferation and viability were assessed using flow cytometry. Bar graphs show fold changes as compared to the Vehicle-treated, non-activated cells. Paired t-test was used for analysis. Each dot represents one donor (n=3 different donors).

Figure S6

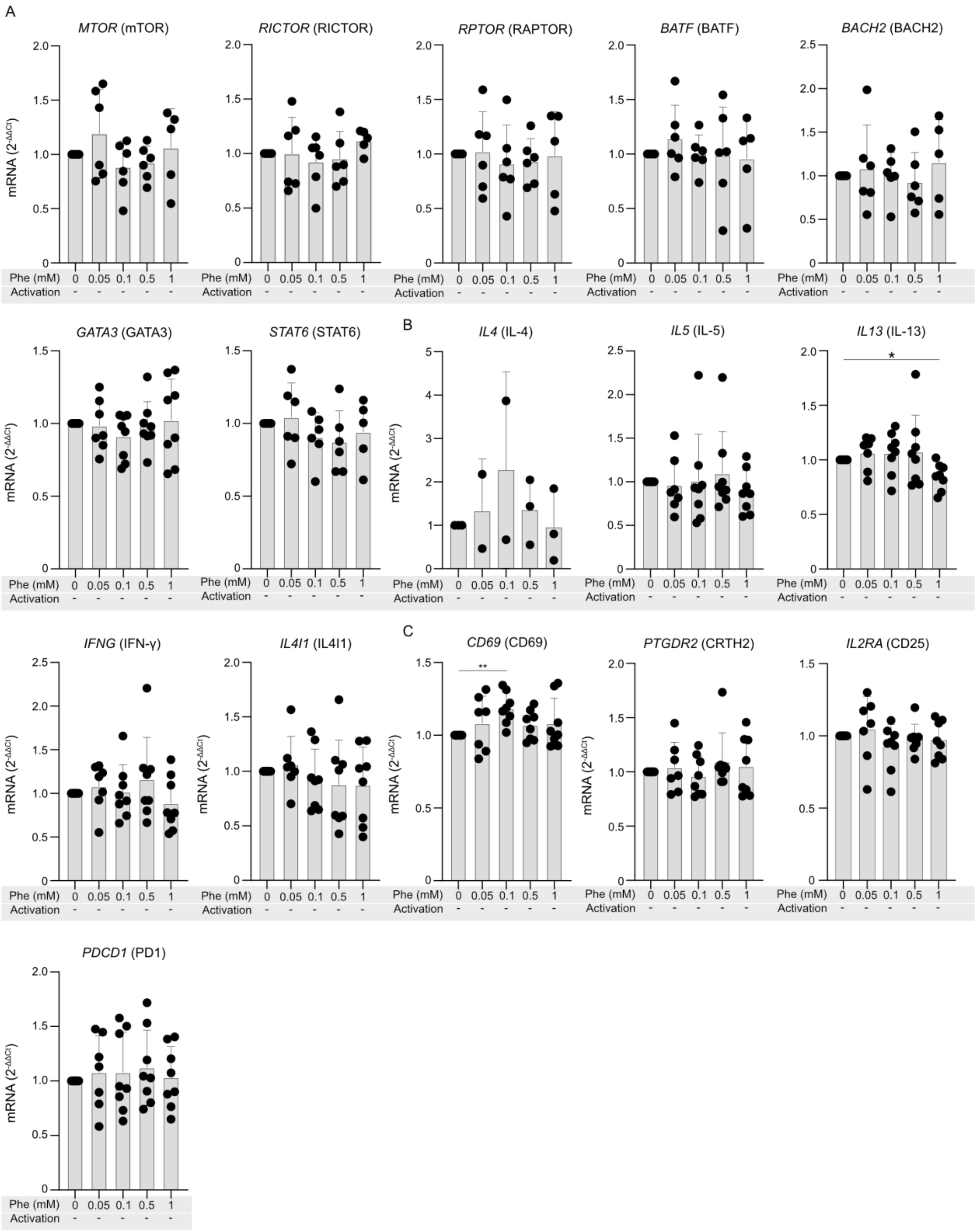


**Figure S6. Effect of L-phenylalanine on the mRNA expression of type 2 transcription factors, cytokines, enzymes, and surface markers in non-activated Th2 cells**. (**A-C**) mRNA expression of transcription factors (**A**), cytokines and enzymes (**B**), and surface markers (**C**) in *in vitro* differentiated Th2 cells treated with increasing doses of Phe. Th2 cells were incubated in full cell culture medium supplemented with Vehicle, 0.05mM, 0.1mM, 0.5mM and 1mM Phe without activation with CD2, CD3 and CD28 antibodies for 24h. Following incubation, total RNA was isolated and mRNA expression was determined using qRT-PCR. Data are analysed using One-way ANOVA with Dunnett's correction (n=6-8 different donors). Bars represent mean±SEM. *p<0.05, **p<0.01, ***p<0.001.

Figure S7

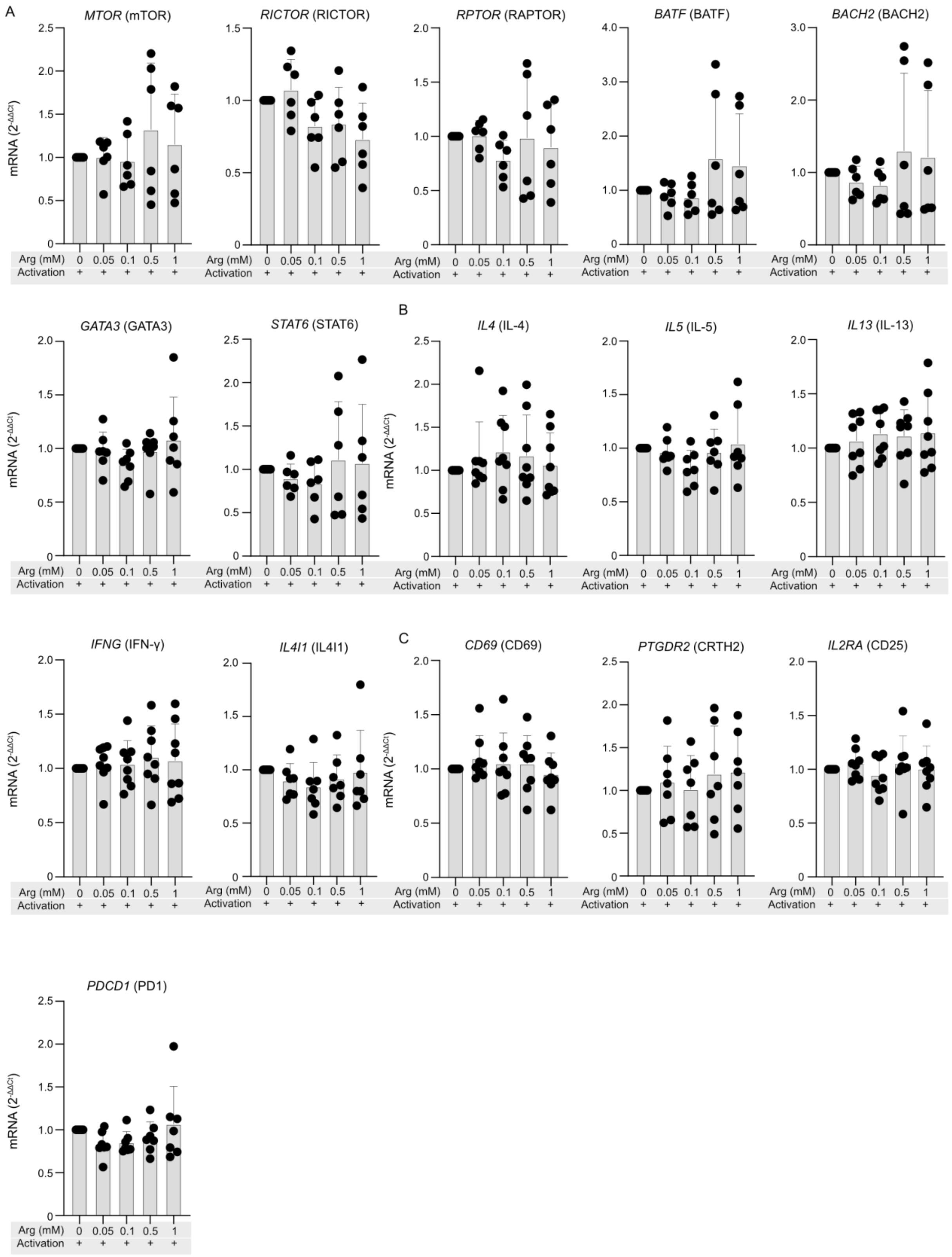


**Figure S7. Effect of L-arginine on the mRNA expression of type 2 transcription factors, cytokines, enzymes and surface markers in activated Th2 cells.** (**A-C**) mRNA expression of transcription factors (**A**), cytokines and enzymes (**B**), and surface markers (**C**) in *in vitro*

differentiated Th2 cells treated with increasing doses of Arg. Th2 cells were incubated in full cell culture medium supplemented with Vehicle, 0.05mM, 0.1mM, 0.5mM and 1mM Arg with activation with CD2, CD3 and CD28 antibodies for 24h. Following incubation, total RNA was isolated and mRNA expression was determined using qRT-PCR. Data are analysed using One-way ANOVA with Dunnett's correction (n=6-8 different donors). Bars represent mean±SEM. *$p<0.05$, **$p<0.01$, ***$p<0.001$.

Figure S8

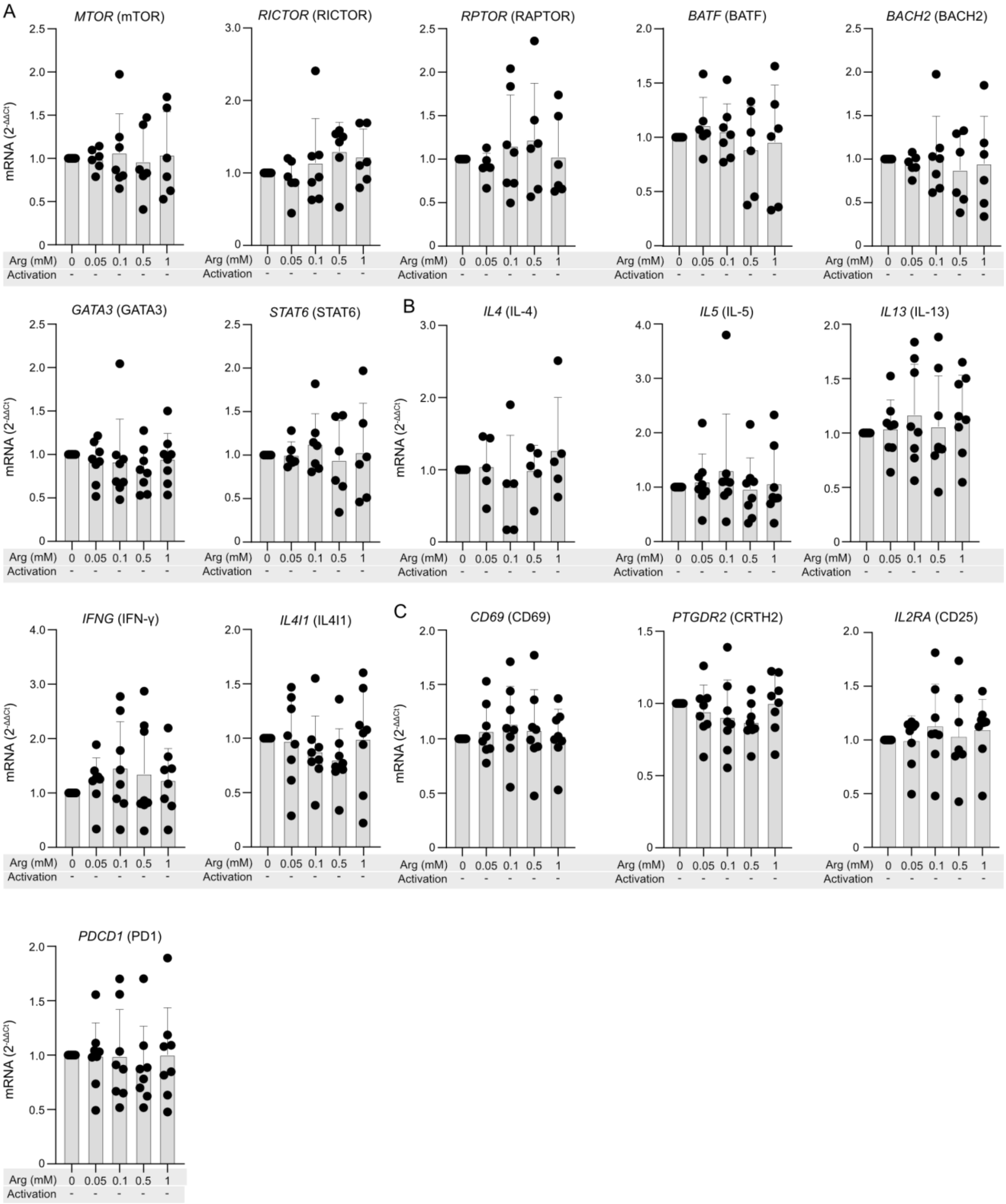


**Figure S8. Effect of L-arginine on the mRNA expression of type 2 transcription factors, cytokines, enzymes, and surface markers in non-activated Th2 cells.** (**A-C**) mRNA expression of transcription factors (**A**), cytokines and enzymes (**B**), and surface markers (**C**) in *in vitro* differentiated Th2 cells treated with increasing doses of Arg. Th2 cells were incubated in full cell culture medium supplemented with Vehicle, 0.05mM, 0.1mM, 0.5mM and 1mM Arg without activation with CD2, CD3 and CD28 antibodies for 24h. Following incubation, total

RNA was isolated and mRNA expression was determined using qRT-PCR. Data are analysed using One-way ANOVA with Dunnett's correction (n=6-8 different donors). Bars represent mean±SEM. *p<0.05, **p<0.01, ***p<0.001.

Figure S9

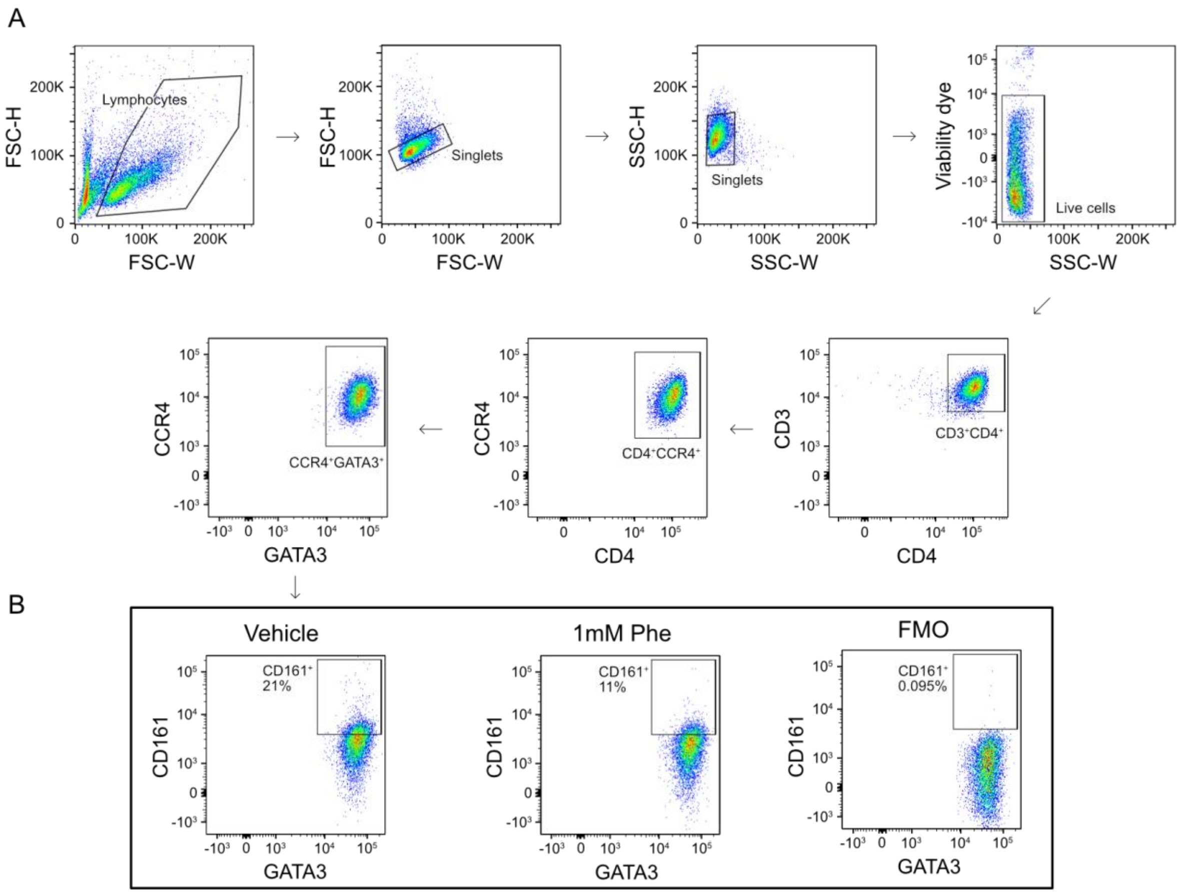


**Figure S9. Effect of L-phenylalanine supplementation on Th2a cells.** (**A**) *In vitro* differentiated Th2 cells were incubated in media with or without supplementation of 1mM Phe for 24h with or without CD2, CD3 and CD28 activation antibodies. Following incubation, they were stained with panel provided in Table S8 and acquired on the flow cytometer. In short, live cells were negatively selected following elimination of doublets. $CD3^+CD4^+$ double positive cells were subsequently gated. Within this population, $CCR4^+$, $GATA3^+$ cells were positively selected and the frequency of $CD161^+$ Th2 (Th2a) cells was assessed in this population. (**B**) Representative flow cytometry plots showing the frequency of $CD161^+$ cells within $CD3^+CD4^+CCR4^+GATA3^+$ Th2 cells from one donor incubated in media additionally supplemented with Vehicle (left) or 1mM Phe (center). FMO control for CD161 signal has been included on the right. Pooled data from 3 different donors are shown in Figure 5F.

Figure S10

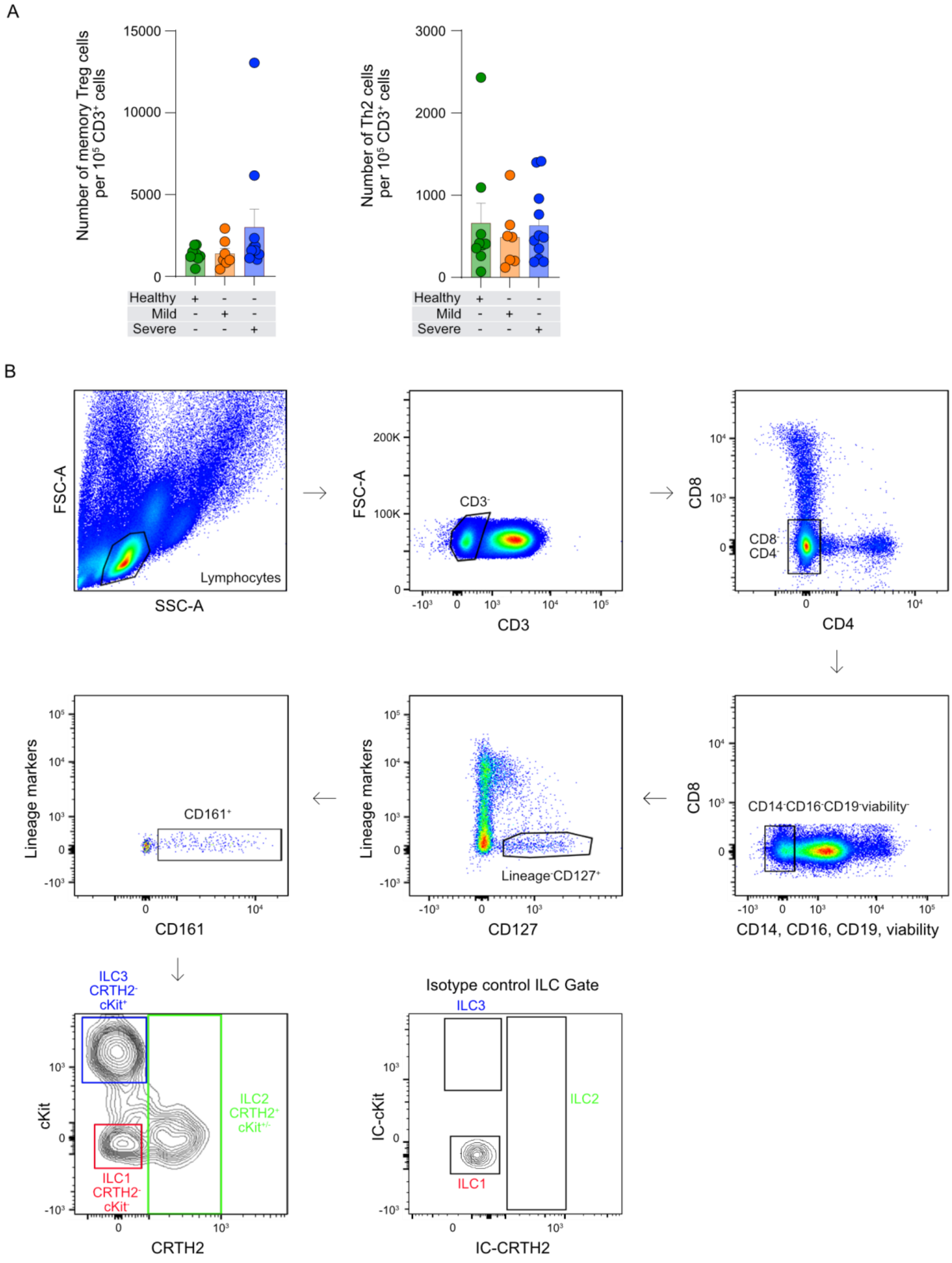


**Figure S10. Flow cytometric assessment of memory cell populations and gating strategy for the study of ILCs in Cohort A samples.** (**A**) Counts of memory Treg cells (left) and total Th2 cells (right) in healthy subjects (n=9), mild-allergic patients (n=7) and severe allergic patients (n=11) from participants in Cohort A. Counts were determined using flow cytometry and analysed with One-way ANOVA with Tukey correction. Bars represent mean±SEM and each dot represents one donor. (**B**) Gating strategy for assessment of innate lymphoid cells (ILCs) followed in classical analysis. ILCs were gated by eliminating doublets and all other immune cell lineages. Within CD127$^+$ and CD161$^+$ cells different classes of ILCs could be discerned by differential expression of CRTH2 and cKit: ILC1 (CRTH2$^-$, cKit$^-$), ILC2 (CRTH2$^+$, cKit$^{+/-}$), and ILC3 (CRTH2$^-$, cKit$^+$).

Figure S11

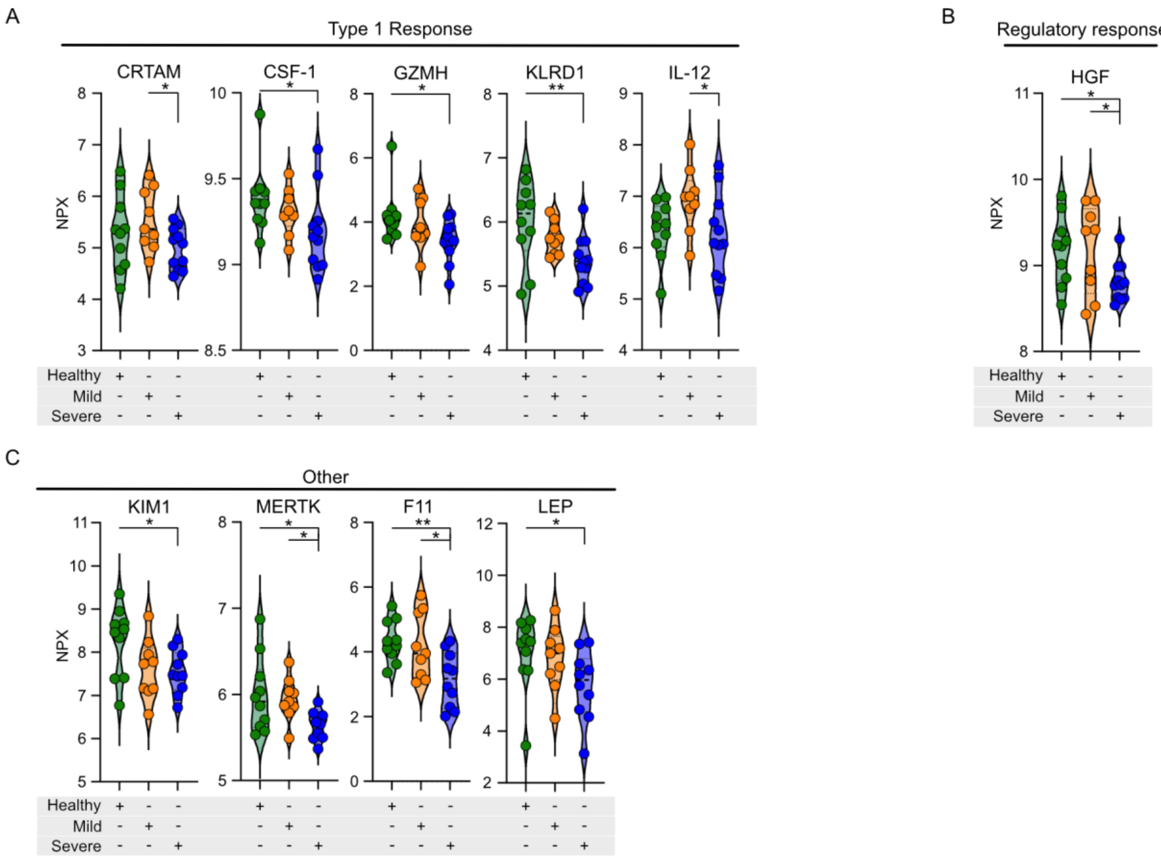


**Figure S11. Systemic type 2 inflammatory profile of subjects from Cohort A by serum proteomics.** (**A-C**) Violin plots of differentially expressed type 1 response (**A**), regulatory response (**B**) and key inflammation associated (**C**) proteins in the serum of controls (n=10), mild (n=9) and severe (n=10) allergic patients (Cohort A) analyzed by Proximity Extension Assay (PEA) and presented as NPX. One-way ANOVA with Fishers LSD test was used to compare differences between groups. All bars represent the mean±SEM and each dot represents one donor. *p<0.05, **p<0.01.

Figure S12

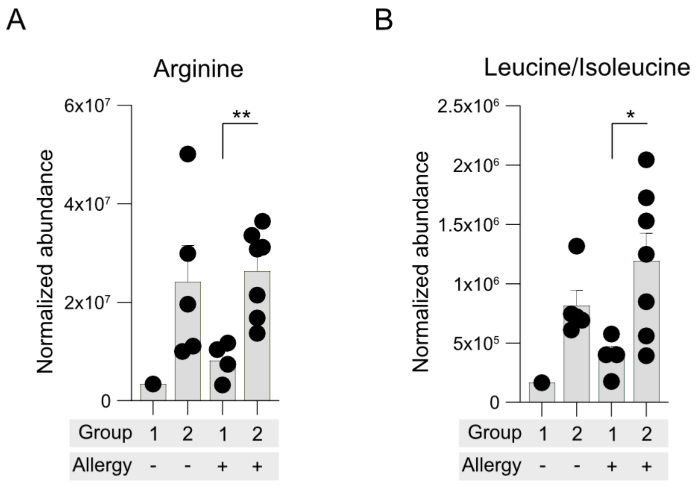


**Figure S12. Quantification of Arg and leucine/isoleucine in Teff cells by metabolomics of Cohort A subjects**. (**A-B**) Abundance of Arg (**A**) and Leu (**B**) in group 1 (control, n = 1; severe allergy, n = 4) and group 2 (control, n = 5; mild allergy = 1, severe allergy, n = 6) (subset of cohort A). Unpaired t-test was used to compare differences between groups. All graphs represent the mean±SEM. *p<0.05, **p<0.01, ***p<0.001.

Figure S13

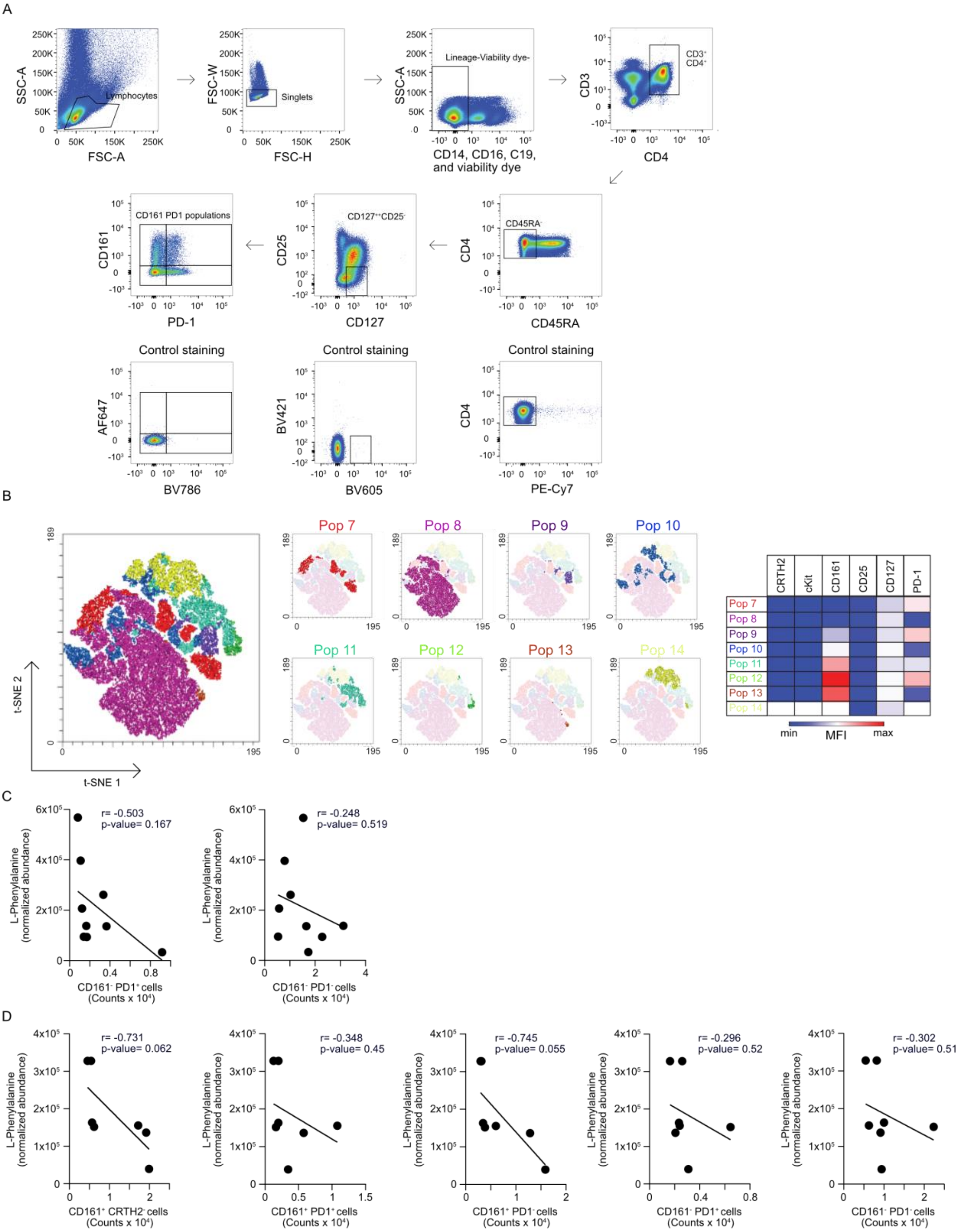


**Figure S13. Unbiased 2-dimensional flow cytometeric analysis of memory CD4$^+$Teff cells from severe allergic patients and healthy controls (subset of Cohort A).** (**A**) Gating strategy used for t-SNE and subsequent classical analysis to determine frequencies of populations based on expression levels of CRTH2, cKit, CD161, and PD-1 by flow cytometric assessment. (**B**) tSNE plot of unbiased 2-dimensional flow cytometric analysis of memory

CD4$^+$Teff cells (CD3$^+$CD4$^+$CD45RA$^-$CD127$^+$CD25$^-$) from healthy controls (subset of Cohort A) identifying seven subpopulations based on the expression levels of CRTH2, cKit, CD161, and PD-1. (**C**) Pearson correlation of normalized abundance of intracellular Phe, measured in memory CD4$^+$Teff cells from patients with severe allergy, with total counts of CD161$^-$ populations within memory CD4$^+$Teff cells (subset of Cohort A). (**D**) Pearson correlation of normalized abundance of intracellular Phe, measured in memory CD4$^+$Teff cells from healthy controls, with total counts of CD161$^+$ and CD161$^-$ populations within memory CD4$^+$Teff cells (subset of Cohort A).

Figure S14

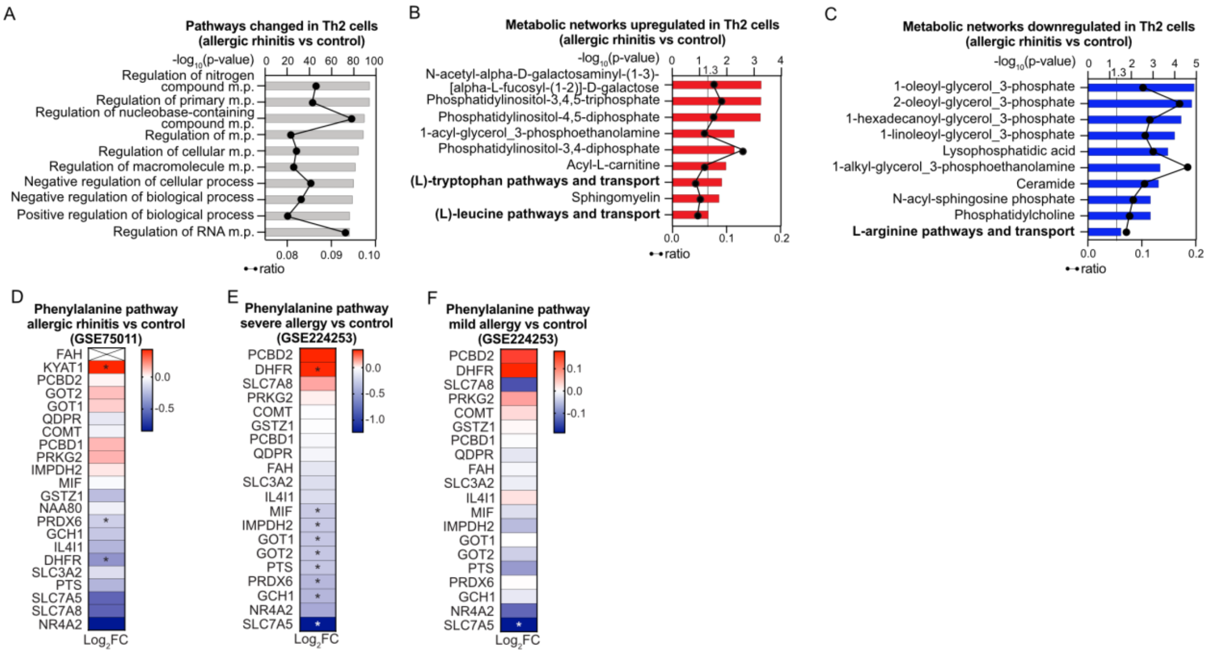


**Figure S14. Expression of L-Phenylalanine metabolism related genes in previously published datasets (Cohorts B and C).** (**A-C**) Top significantly enriched pathways (**A**), significantly upregulated (**B**) and significantly downregulated (**C**) metabolic networks within differentially expressed genes (DEG, $p<0.05$) in allergic rhinitis patients compared to controls (control n=15, allergic rhinitis n=25) from GSE75011[68] (Cohort B). Black line represents ratio of genes in experiment over complete pathway set. All data available in tables S14, S16 and S18, respectively. (**D**) Phe metabolism and transport pathway heatmap showing fold change ($log_2$FC) of differentially expressed genes in allergic rhinitis patients (n=25) compared to controls (n=15) from GSE75011[68] (Cohort B). (**E**) Significant differentially expressed genes related to Phe metabolism in severe allergy patients (n=7) in comparison to healthy controls (n=8) from GSE224253[46] (Cohort C). (**F**) Significant differentially expressed genes related to Phe metabolism in mild allergy patients (n=9) in comparison to healthy controls (n=8) from GSE224253[46] (Cohort C). (B-F) Upregulated and downregulated genes are shown in red and blue, respectively. (D-F) Pathway curated and adapted from GSEA and MSigDB Database.

Figure S15

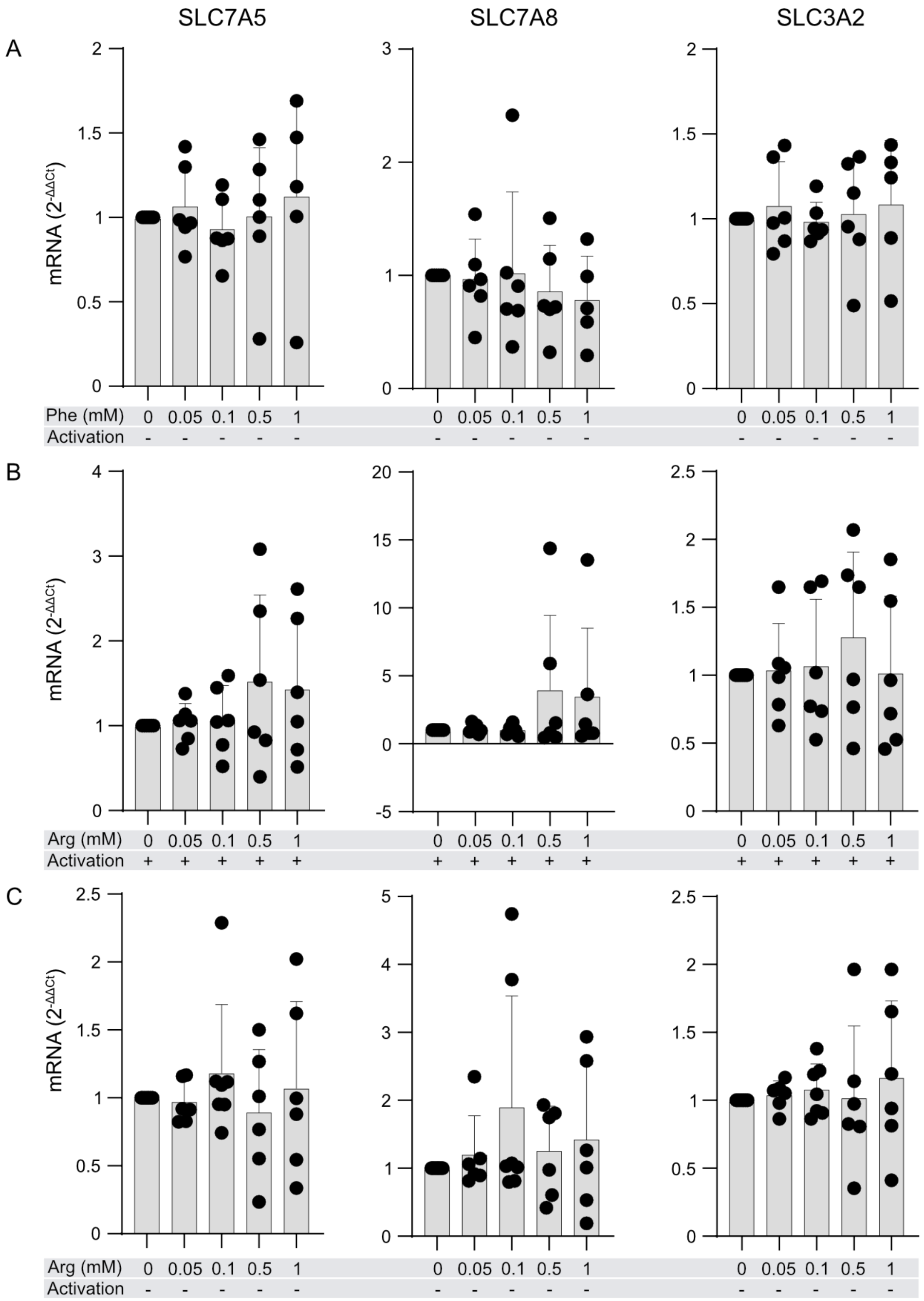


**Figure S15. Effect of L-phenylalanine and L-arginine on the mRNA expression of *SLC7A5* (LAT1), *SLC7A8* (LAT2) and *SLC3A2* (CD98) in Th2 cells.** (A-C) *In vitro* differentiated Th2 cells were treated with additional Phe without CD2, CD3 and CD28 activation (**A**) and additional Arg with (**B**) or without (**C**) CD2, CD3 and CD28 activation for 24h. Following incubation, mRNA expression was determined using qRT-PCR (n=6-8 different donors). Data are analysed using One-way ANOVA with Dunnett's correction (n=6-8 different donors). Bars represent mean±SEM. *p<0.05, **p<0.01, ***p<0.001.

### Supplementary Table Legends

**Table S1.** Antibody panel used for *ex vivo* fluorescence-activated cell sorting and immunophenotyping.

**Table S2.** Summary of identified metabolites. [a]ID number was assigned to metabolites and is shared throughout the tables. [b]Annotations were given according to their m/z and RT values using different annotation methods. When multiple identities were possible, they are separated with / when they are isomers (same formula and adduct, but different compound name), and with // when they are different compounds (different adduct, formula, mass and/or mass error). Possible adducts, masses, errors and formulas are also separated with //. # marks those metabolites that were more interesting for further analyses. If those metabolites had at least 2 entries for the same ID, those with better analytical conditions (most stable adduct, better extraction method, better ionization mode, more confident identification) were marked. [c]Annotation tools used include CEU-mass mediator online tool (CEUMM), Lipid Annotator software (LipidAnnotator), MS-DIAL online database (MS-DIAL), and tandem mass spectrometry experiments (MS2). [d]Biochemical class and subclass followed Human Metabolome Database (HMDB) classification. [e]Cell subtype in which the metabolite was found, considering both healthy and allergic subjects. [f]Relative Standard Deviation (or Coefficient of variation, CV, in %) in the QC samples for both Treg and Teff QCs. [g]Reason why the metabolite was excluded from the Treg or Teff subset: '% Blanks': if the average signal of each feature in the blanks was higher than the average in the samples; '%CV QC>30%': if CV_QC>30% for either Teff or Treg; 'Blank subtraction': the average signal of each feature in the blanks was subtracted to its signal in each QC and sample, and only those features where it remained positive in at least 75% of the samples or in the QCs were maintained; and 'Correlation Calibration Curve': if the feature did not have a significant (p-value<0.05) correlation with a correlation coefficient (|ρ|)>0.6. Metabolites in red were only found in allergic patients. [&]marks one metabolite that was not found in non-allergic patients for Treg; although it did pass quality criteria for both cell subpopulations considering allergic patients. Legend: m/z: mass-to-charge ratio; RT: Retention Time; QC: Quality Control; CV: Coefficient of Variation.

**Table S3.** Biochemical composition of the metabolome of memory Teff and Treg cells, according to HMDB database, based on total metabolites. N/A: not available. CAR: carnitine; LPA: lysophosphatidic acid; AMPc: adenosine monophosphate cyclic; IMPc: inosine monophosphate cyclic; SM: sphingomyelin; dUTP: deoxy uracyl triphosphate; UMPc: uracyl monophosphate cyclic.

**Table S4.** Unbiased overrepresentation pathway analysis based on shared (between memory Teff and memory Treg) and memory Teff unique metabolites. List of pathways from KEGG database found in the IMPaLA analysis for Teff cells. Significant pathways (p-value<0.05) are shown in bold, whereas those included in Figure 1G are highlighted in yellow and italics. [a]Number of metabolites found in the pathway from the total number (in this case, 48 metabolites) included in the analysis. [b]Total number of metabolites in the pathway, with the number found in the background list in brackets. Legend: FDR: False Discovery Rate.

**Table S5.** Unbiased overrepresentation analysis based on shared (between memory Teff and memory Treg) and memory Treg unique metabolites. List of pathways from KEGG database found in the IMPaLA analysis for Treg cells. Significant pathways (p-value<0.05) are shown in bold, whereas those included in Figure 1G are highlighted in yellow and italics. [a]Number of metabolites found in the pathway from the total number (in this case, 51 metabolites) included in the analysis. [b]Total number of metabolites in the pathway, with the number found in the background list in brackets. Legend: FDR: False Discovery Rate.

**Table S6.** Biochemical composition of the metabolome of Teff and Treg cells according to HMDB database based on shared features only. N/A: not available. CAR: carnitine; LPA: lysophosphatidic acid; AMPc: adenosine monophosphate cyclic; dUTP: deoxy uracyl triphosphate.

**Table S7.** Pairwise comparisons of features present in both Teff and Treg subsets. For exploratory purposes, we considered metabolites with a p-value<0.05 for the following analyses. # marks those metabolites that were more interesting for further analyses. If those metabolites had at least 2 entries for the same ID, those with better analytical conditions (most stable adduct, better extraction method, better ionization mode, more confident identification) were marked. Legend: m/z: mass-to-charge ratio; RT: Retention Time; FDR: False Discovery Rate.

**Table S8.** Antibody panel used for confirmation of Th2 status of *in vitro* differentiated Th2 cells and Single Cell ENergetIc metabolism by profilIng Translation inHibition (SCENITH).

**Table S9.** Demographic of all recruited participants in Cohort A. For categorical and continous variables, the chi square test and Kruskal Wallis with Dunn's test for multiple comparisons, respectively, were used. Significance was set as *p<0.05; **p<0.01; ***p<0.001. F: Female; IgE: Immunoglobulin E; kU/L: kilounit per liter, SD: Standard deviation; FEV1: Forced expiratory volume; FVC: Forced vital capacity; avg: average; N/A: not applicable.

**Table S10.** Expanded demographics of allergic patients and healthy controls recruited in the study as part of Cohort A. F: Female; M: Male; IgE: Immunoglobulin E; kU/L: kilounit per liter;

Ole e: *Olea europaea*; rPhlp: recombinant *Phleum pratense*; Der p: *Dermatophagoides pteronyssinus*; rPol d: recombinant *Polistes dominulus*; Alt a: *Alternaria alternata*; n Sal k: natural *Salsola kali*; ND: not determined/not available.

**Table S11.** Pairwise comparisons of features present in the memory $CD4^+$Teff subset between allergic and non-allergic individuals. Statistical significance was set for FDR < 0.05. # marks those metabolites that were more interesting for further analyses. If those metabolites had at least 2 entries for the same ID, those with better analytical conditions (most stable adduct, better extraction method, better ionization mode, more confident identification) were marked. Metabolites in red were only found in allergic patients. [&]marks one metabolite that was not found in non-allergic patients for Treg; although it did pass quality criteria for both cell subpopulations considering allergic patients. Legend: m/z: mass-to-charge ratio; RT: Retention time; FDR: False Discovery Rate.

**Table S12.** Pairwise comparisons of features present in the memory Treg subset between allergic and non-allergic individuals. Statistical significance was set for FDR < 0.05. # marks those metabolites that were more interesting for further analyses. If those metabolites had at least 2 entries for the same ID, those with better analytical conditions (most stable adduct, better extraction method, better ionization mode, more confident identification) were marked. Metabolites in red were only found in allergic patients. [&]marks one metabolite that was not found in non-allergic patients for Treg; although it did pass quality criteria for both cell subpopulations considering allergic patients. Legend: m/z: mass-to-charge ratio; RT: Retention time; FDR: False Discovery Rate.

**Table S13.** Enrichment analysis of the most significant process networks using all significantly changed genes in Th2 cells in allergic asthma patients compared to healthy controls in Cohort B. Legend: FDR: False Discovery Rate.

**Table S14.** Enrichment analysis of the most significant process networks using all significantly changed genes in Th2 cells in allergic rhinitis patients compared to healthy controls in Cohort B. Legend: FDR: False Discovery Rate.

**Table S15.** Enrichment analysis of significantly upregulated metabolic networks in Th2 cells in allergic asthmatic patients compared to healthy controls in Cohort B using significantly upregulated genes. FDR: False Discovery Rate.

**Table S16.** Enrichment analysis of significantly upregulated metabolic networks in Th2 cells in allergic rhinitis patients compared to healthy controls in Cohort B using significantly upregulated genes. FDR: False Discovery Rate.

**Table S17.** Enrichment analysis of significantly downregulated metabolic networks in Th2 cells in allergic asthmatic patients compared to healthy controls in Cohort B using significantly downregulated genes. FDR: False Discovery Rate.

**Table S18.** Enrichment analysis of significantly downregulated metabolic networks in Th2 cells in allergic rhinitis patients compared to healthy controls in Cohort B using significantly downregulated genes. FDR: False Discovery Rate.

**Table S19.** Curated and adapted gene set for Phenylalanine pathway analysis from GSEA and MSigDB Database (Broad Institute, MIT and Reagent of the University of California, USA) available under systematic names: M29207, M37685, M46134, M16650, M27835, M27850, M45537 used in the assessment of previously published transcriptomic data.

**Table S20.** Demographics of allergic patients and healthy controls in Cohort D. Legend: F: Female; M: Male; GINA: Global Initiative for asthma classification; ACT: asthma control test; MA: mild allergic; SA: severe allergic; Ole e: *Olea europaea*; kU/L: kilounit per liter; N/A: not applicable; N/D: not determined. No patients were hospitalized.

**Table S21.** Demographics of healthy subjects and allergic patients in Cohort E. Legend: F: Female; M: Male; BMI: Body-Mass-Index; IgE: Immunoglobulin E; kU/L: kilounit per liter; Bet v: *Betula verrucosa*; Phl p: *Phleum pratense*; Ole e: *Olea europaea*; FEV1: Forced expiratory volume.
